# Integrated phylogenomic analyses reveal recurrent ancestral large-scale duplication events in mosses

**DOI:** 10.1101/603191

**Authors:** Bei Gao, Moxian Chen, Xiaoshuang Li, Yuqing Liang, Daoyuan Zhang, Andrew J. Wood, Melvin J. Oliver, Jianhua Zhang

## Abstract

- Mosses (Bryophyta) are a key group occupying important phylogenetic position for understanding land plant (embryophyte) evolution. The class Bryopsida represents the most diversified lineage and contains more than 95% of the modern mosses, whereas the other classes are by nature species-poor. The phylogeny of mosses remains elusive at present.
- Recurrent whole genome duplications have shaped the evolution trajectory of angiosperms, but little is known about the genome evolutionary history in mosses. It remains to be answered if there existed a historical genome duplication event associated with the species radiation of class Bryopsida.
- Here, the high-confidence moss phylogeny was generated covering major moss lineages. Two episodes of ancient genomic duplication events were elucidated by phylogenomic analyses, one in the ancestry of all mosses and another before the separation of the Bryopsida, Polytrichopsida and Tetraphidopsida, with estimated ages of the gene duplications clustered around 329 and 182 million year ago, respectively.
- The third episode of polyploidy event (termed ψ) was tightly associated with the early diversification of Bryopsida with an estimated age of ~87 million years. By scrutinizing the phylogenetic timing of duplicated syntelogs in *Physcomitrella patens*, the WGD1 and WGD2 events were confidently re-recognized as the ψ event and the Funarioideae duplication event (~65 mya), respectively. Together, our findings unveiled four episodes of polyploidy events in the evolutionary past of *Physcomitrella patens*.

## Introduction

Gene duplications provided the raw genetic materials for evolutionary innovations and are considered as important driving forces in species diversification and evolution (Ohno, 1970; Van de Peer *et al.*, 2017) and the evolutionary past of land plants is characterized by recurrent ancestral polyploidy events (Blanc & Wolfe, 2004; Cui *et al.*, 2006; Van de Peer *et al.*, 2017). The duplicated genes in *Arabidopsis* can be traced back to five rounds of polyploidy termed the α-β-γ-ε-ξ series dating back to the origin of seed plants, where the ε event occurred in the ancestor of angiosperms and ξ for all seed plants (Blanc *et al.*, 2003; Bowers *et al.*, 2003; Jiao *et al.*, 2011), though these two ancestral polyploidy events are still under debate and questioned in a recent study (Ruprecht *et al.*, 2017). Echoing the α-β-γ series in the *Arabidopsis thaliana* lineage, a WGD series of ρ-σ-τ: were reported in rice and other grasses (Jiao *et al.*, 2014). Notably, the gamma event has been extensively studied and systematic molecular dating results suggest this ancestral genome triplication event clustered in time with the early diversification of core-eudicots in Aptian of late Cretaceous. The gamma event thus appears to be associated with the species radiation of more than 75% of extant angiosperms which contributed to answer the Darwin’s “abominable mystery” (Friedman, 2009; Jiao *et al.*, 2012; Vekemans *et al.*, 2012). However, both phylogenetic relationships and the evolutionary history of mosses, one of the most important monophyletic clade in the early-diverging land plant lineages (Morris *et al.*, 2018) comprising more than twelve thousand species (Goffinet, 2004; Shaw *et al.*, 2011), remains largely uncharacterized.

In an early study utilizing a collection of transcript datasets, a polyploidy event was identified in the evolutionary past of the model moss *Physcomitrella patens* (Rensing *et al.*, 2007) and with the completion of the *P. patens* genome the polyploidy event was characterized by a *Ks* peak around 0.5-0.9 (Rensing *et al.*, 2008). A more recent report describing the chromosome-scale assembly of the *P. patens* genome further deciphered the large-scale duplication event as two recurrent WGDs: the younger WGD2 with a *Ks* peak around 0.5-0.65 and an older WGD1 with a *Ks* peak around 0.75-0.9 (Lang *et al.*, 2018). In a supplementary analysis, paralogous *Ks* frequency distribution in a number of moss transcriptomes were also analyzed to indicate the widespread ancestral genomic duplications in mosses (Lang *et al.*, 2018). The detailed statistical analyses of the *Ks* distribution of paralogs in *Ceratodon purpureus* transcriptome also suggested multiple ancestral genomic duplications (Szovenyi *et al.*, 2015) and a phylogenomic study reported multiple genome duplications in peat mosses (class Sphagnopsida), as well as one ancestral polyploidy event that occurred in the ancestor of mosses (Devos *et al.*, 2016).

Advances in high-throughput sequencing technologies have generated adequate transcriptome data for the seeking of signals of ancestral large-scale genome duplications by analyzing the Ks-based age distributions (Blanc & Wolfe, 2004; Cui *et al.*, 2006) and/or using large-scale gene tree-based phylogenomic approaches (Jiao *et al.*, 2011; Jiao *et al.*, 2012; Yang *et al.*, 2018). For example, the 1000 Green Plant Transcriptome Project (1KP, https://www.onekp.com) generated at least three Gb of paired-end Illumina reads for one thousand plants covering most lineages from green algae to angiosperms. In the OneKP pilot study, transcriptome data for a total of 92 plants were generated and among them 11 moss species were included (Wickett *et al.*, 2014). The transcriptomes in two desert mosses *Syntnchia caninervis* and *Bryum argenteum* were reported in our previous studies (Gao *et al.*, 2014; Gao *et al.*, 2017) and a recent phylotranscriptomic study in pleurocarpous mosses (Hypnales) generated a large set of transcriptomic data (Johnson *et al.*, 2016).

Despite the rapid growing body of the sequencing data in mosses, the evolutionary past, especially the large-scale genome duplication events and the precise phylogenetic positioning of these events remain unresolved in the Bryopsida, the largest class of mosses containing more than 95% of the extant moss species (Newton *et al.*, 2006). The detection of these ancestral duplication events has been hampered by the undetermined phylogenetic relationships of the Bryopsida with other classes of mosses, although some taxonomic and phylogenetic relationships have been proposed (Shaw *et al.*, 2010a; Shaw *et al.*, 2010b; Shaw *et al.*, 2011).

The *P. patens* chromosome structure and intra-genomic synteny comparison provided unequivocal evidence for two recurrent ancestral WGDs (Lang *et al.*, 2018), but the precise phylogenetic position for the two events was unresolved. Whether or not the WGDs were shared with other mosses or simply species-specific polyploidy events remain an open question. Although inter-genome synteny comparison remains unavailable at present because of the lack of a closely related moss genome, resolving the phylogenetic placement of the two *P. patens* WGDs could be achieved by investigating the phylogenetic timing of the duplicated syntelog pairs. For example, the precise phylogenetic timing of the core-eudicot gamma triplication event was successfully resolved by tracing and summarizing the phylogenetic timing of *Vitis* syntelogs (Jiao *et al.*, 2012).

To resolve the phylogeny of the mosses and to provide the robust evidence and probe the phylogenetic position for ancestral genomic duplication events, we have utilized phylotranscriptomic analyses utilizing augmented transcriptomic sequence data, a proven robust methodology to generate reliable species phylogenies (Wickett *et al.*, 2014; Ran *et al.*, 2018; Unruh *et al.*, 2018). This was coupled with a large-scale phylogenomic analyses investigating duplication signals from thousands of gene-trees to determine the phylogenetic positioning for ancestral genomic duplication events and that avoids the limitations of paranome-based *Ks* frequency studies (Jiao *et al.*, 2011; Devos *et al.*, 2016; Johnson *et al.*, 2016).

We compiled transcriptomic data from diversity of moss lineages from either publicly available collections or *de novo* assembled from deposited transcript sequence reads. A total of twenty-seven moss transcriptomes were included in our phylogenomic analyses representing six distinctive classes (20 in Bryopsida, 2 Polytrichopsida, 1 Tetraphidopsida, 1 Andreaeopsida, 2 Sphagnopsida and 1 Takakiopsida) in Bryophyta. The transcriptomes (species) to include in the analyses were chosen to enable the division of long internal branches in the phylogeny that may lead to artifacts in the analyses (Jiao *et al.*, 2012) and that generate largely laddered species phylogenies (Li *et al.*, 2015). We included the gene sequences from the genomes of *Physcomitrella patens* along with those of the liverwort (*Marchantia polymorpha*) and the early-diverging tracheophyte (*Selaginella moellendorffii*), the latter two genomic collections were used to derive the correct rooting of the gene trees and to provide informative nodes for molecular dating. The transcriptomic and genomic data were sorted into gene families, followed by a phylogenomic cleaning and masking procedure (Yang & Smith, 2014). One-to-one orthogroups were collected and employed to reconstruct the moss phylogeny, while maximum-likelihood gene trees for orthogroups containing multi-copy genes were utilized for the identification of ancestral duplication events. Based on these gene trees, the phylogenetic timing of the *P. patens* paralogous pairs retained in genomic synteny regions were investigated and traced to identify the phylogenetic positioning of the two WGD events.

## Materials and Methods

### Transcriptome data collection

Transcriptomes or genomes from a total of 29 land plant species, including one lycophyte (*Selaginella moellendorffii*), one liverwort (*Marchantia polymorpha*), and 27 from Bryophyta (mosses) that covered a wide range of moss lineages were collected. A detailed list of species and corresponding lineages are presented in Table S1. To optimize computational capacity we limited the analysis to moss species that occupy significant phylogenetic positions in the moss phylogeny rather than include all currently available moss transcriptomes (Wickett *et al.*, 2014; Johnson *et al.*, 2016; Lang *et al.*, 2018). A large number of transcriptomes in Hypnales were generated (Johnson *et al.*, 2016), but only five were used in this analysis as representatives from different phylogenetic clades. For the subclass Dicranidae, reads or transcriptome sequences were included for five species from four studies (Gao *et al.*, 2014; Wickett *et al.*, 2014; Johnson *et al.*, 2016; Lang *et al.*, 2018). If assembled transcriptomes were not available, the Illumina reads deposited in the NCBI-SRA were obtained, cleaned using Trimmomatic v0.36 (Bolger *et al.*, 2014), and *de novo* assemblies were generated using Trinity v2.6.6 (Haas *et al.*, 2013). For all the transcript sequences generated, coding region sequences were predicted and translated into peptide sequences using the TranDecoder v5.3.0 pipeline (https://github.com/TransDecoder/), and the cd-hit v4.6 algorithm (Li & Godzik, 2006) was employed to remove redundantly assembled sequences with sequence identity threshold of 0.98 (-c 0.98) and minimum sequence coverage of 0.9 (-aS 0.9). Only peptide sequences with a minimum length of 50 amino acids were retained for further analyses. Coding sequences derived from the genomes of the model plant species (*Physcomitrella patens, Marchantia polymorpha*, and *Selaginella moellendorffii)* were downloaded from Phytozome v12.1.6 database (https://phytozome.jgi.doe.gov/).

### Clustering transcripts into orthologous groups

To cluster transcripts into orthologous groups, all-by-all pair-wise bidirectional blastp (Buchfink *et al.*, 2015) comparisons (e-value ≤ 1e-5) were conducted among the 29 transcriptomes, and only alignments with greater than 50% coverage were retained and introduced into the OrthoFinder v2.2.6 pipeline (Emms & Kelly, 2015) for orthologous group classification. Orthogroups were required to have at least one sequence from each of the three model plant genomes (*P. patens, M. polymorpha* and *S. moellendorffii*) and orthogroups with less than 10 sampled taxa were discarded; a total of 5,684 orthogroups were retained for further analyses.

Protein sequences in each orthologous cluster that represent no less than 10 taxa were aligned using MAFFT v7.310 (Katoh & Standley, 2013) employing the L-INS-i strategy and a maximum of 5000 iterations. Coding sequences were forced onto the protein sequence alignment using Phyutility v2.2.6 and trimmed by a minimal column occupancy of 0.1 (Smith & Dunn, 2008). Gene trees for each orthologous group were constructed using FastTree v2.1.10 (Price *et al.*, 2010) with 1000 bootstrap replicates. At this stage in the phylogenomic pipeline (Fig. S1), gene clusters may include alternatively spliced isoforms assembled by Trinity and/or paralogs with high sequence identity. The Yang & Smith pipeline (Yang & Smith, 2014) was employed to inspect monophyletic or paraphyletic clades in each of the trees consisting of genes from a single taxon, and these clades were reduced to retain only sequences with the longest unambiguous alignment. In this way, both the alternatively spliced transcript isoforms and recently duplicated paralogs in each transcriptome were masked after the Yang & Smith pipeline.

After gene tree-based cleaning and masking, the original protein sequences in each gene cluster (before trimming) were realigned using MAFFT v7.310 (Katoh & Standley, 2013). The protein alignments were trimmed (-automated1) and back-translated using TRIMAL v1.4 (Capella-Gutierrez *et al.*, 2009). The generated coding sequence alignments for orthogroups were divided into two subsets: one subset containing the putatively one-to-one orthogroups (in such orthogroups each taxa contains no more than one gene sequence) and was employed in the species tree estimation and the other subset, containing multi-copy orthogroups, was used to identify ancestral gene duplications in mosses.

### Species tree construction

The generated orthogroups were masked to maximize the number of orthogroups where each species was represented by a single transcript (Table S2). Such a subset of putatively deduced one-to-one orthologs or single-copy gene families were utilized for species tree estimation using both concatenated sequence supermatrix and coalescence-based summary methods (Johnson *et al.*, 2016; Unruh *et al.*, 2018). But it must be acknowledged that even in these putatively single-copy gene families, some ancient paralogs may be misidentified as orthologs due to lineage or species-specific non-orthologous gene losses (Robertson *et al.*, 2017). Reduced coding sequence alignments for the 649 single-copy orthogroups were concatenated, and gaps inserted where taxa were missing. Three partition strategies were employed for a concatenated alignment: (I) original concatenated alignment unpartitioned; (II) one partition for the first and second codon positions, and a separate partition for the third codon position; (III) the partition scheme recommended by PartitionFinder v2.1.1 (Lanfear *et al.*, 2017) that treated each codon position in each gene as individual partitions, and the best-fit partitioning scheme divided concatenated alignment into 611 partitions. The concatenated species tree was then estimated using RAxML v8.2.12 (Stamatakis, 2014) under the GTRGAMMA evolutionary model and 500 bootstrap replicates with *Selaginella moellendorffii* set as the outgroup. To further validate the species tree, we also constructed species trees using the first and second codon positions and the concatenated protein alignments under the PROTGAMMAAUTO model.

For the coalescence-based summary approach, individual maximum-likelihood unrooted gene trees for CDS alignments were estimated for each of the 649 single-copy orthogroups using RAxML v8.2.12 (Stamatakis, 2014) under the GTRGAMMA model with 500 bootstrap replicates. ASTRAL-III v5.6.2 algorithm (Zhang *et al.*, 2018) was used to estimate the coalescence-based species tree from the individual gene trees, and the support values were calculated by performing 100 replicates of multi-locus bootstrapping.

### Identification the phylogenetic placement of paleopolyploidy events

The 5,035 gene clusters not found in the one-to-one families were employed for the identification of ancestral large-scale gene duplications. Maximum-likelihood gene trees were constructed for each orthogroup using RAxML v8.2.12 under the GTRGAMMA model with 100 rapid bootstraps and with the gene sequences from *Selaginella* or *Marchantia* (when *Selaginella* sequence not available in the orthogroup) as outgroups.

The Phylogenetic Placement of Polyploidy Using the Genomes (PUG) algorithm (McKain *et al.*, 2016; Unruh *et al.*, 2018) was employed to identify paleopolyploidy events using the species tree generated from the concatenated sequence supermatrix as a guide. The PUG algorithm was run for each of the 5,035 gene trees using the “estimate_paralogs” option to count the number of unique duplication nodes. For each potential paralog pair, the coalescence node (potential duplication node) was identified from the gene family phylogeny and species composition for both the duplication clades (subtrees) and the sister lineage were queried and scrutinized to count the duplication events for the internal nodes in the species tree. Only if the combined species composition of the paralog subtree and the sister lineage resolve to the same node in the species tree was the placement considered acceptable for the duplication of that putative paralog pair (McKain *et al.*, 2016; Unruh *et al.*, 2018). Results from PUG were filtered for a minimal bootstrap value of 50 and 80 for each paralog pair coalescence node in gene trees of each orthologous group.

Paralogous pairs retained on syntenic blocks (syntelogs) in the *Physcomitrella patens* genome were identified using the McScanX algorithm (Tang *et al.*, 2008; Wang *et al.*, 2012) and their phylogenetic timing were traced from corresponding gene family phylogenies. A similar strategy was employed previously to analyze the core-eudicot GAMMA event in eudicots by tracing the phylogenetic positions of *Vitis* syntelogs (Jiao *et al.*, 2012).

### Molecular dating of ancestral duplications

The best maximum-likelihood gene trees constructed by RAxML for each orthogroup were used to estimate the age of the duplication nodes using the r8s software package (Sanderson, 2003). The semi-parametric penalized likelihood approach (method=PL) employing a Truncated Newton optimization algorithm (algorithm=TN) were used for gene duplication age estimations. The optimum smoothing parameters for each tree were selected by running a prior r8s cross-validation procedure. Because of the scarcity of reliable fossil records in mosses, we employed two ancestral age constraints in our estimation procedure derived from values reported in the updated report (Morris *et al.*, 2018): a minimum age 474 mya and a maximum age of 515 mya for the ancestral node of the land plants and a fixed constraint age of 424 mya for the ancestor of mosses and liverwort in the gene trees. The fixed constraint age was chosen from the mean age estimations of 443.6-405.3 mya (assuming the Bryophyta clade is monophyletic) or 442.0-405.3 mya (assuming the Hornworts are sister to other land plants) (Morris *et al.*, 2018), as the r8s package requires at least one internal node in the tree to be fixed. These constraints allowed for the estimation of the divergency times of the gene duplications in mosses without further age constraints, and if the age of the duplications at focal nodes were clustered in geological time, the ancestral large-scale duplications in mosses could be circumscribed with a high degree of confidence (Jiao *et al.*, 2011). The inferred duplication nodal ages were collected and then analyzed by EMMIX (McLachlan *et al.*, 1999) with one to ten components, using 100 random and 10 k-means starting values. The Bayesian Information Criterion (BIC) was used to select the best number of components fit to the data.

### Synonymous divergency estimation of duplicated genes

Divergence patterns for duplicated genes were explored by calculating the synonymous substitutions per synonymous site (*Ks*, also known as *Ds)* for each paralogous pair. To capture the ‘pure’ *Ks* peak signal of ancestral duplications, the paralogous pairs with a coalescence node at focal duplication nodes were extracted from the gene trees with a minimum bootstrap value (BSV) of 50%. Protein sequence alignments were generated (Edgar, 2004) and back-translated (Suyama *et al.*, 2006) for each duplicated sequence pair, then alignment gaps and stop codons were removed. *Ks* values for each paralogous gene pair were calculated using KaKs_Calculator v2.0 software (Wang *et al.*, 2010) with the ‘‘GY” model (Goldman & Yang, 1994). The derived paralogous *Ks* values in each species were further analyzed using EMMIX (McLachlan *et al.*, 1999) as described above.

## Results and Discussion

### Reduced transcriptomes and orthogroups

A total of 29 species (Fig. 1) were included for the phylogenomic analyses, including 12 *de novo* assembled transcriptomes generated from Illumina reads (Table S1). The size of the non-redundant protein sequences for each species ranged from 11,140 (*Rhodobryum ontariense*) to 45,329 (*Eosphagnum inretortum*). All counts of filtered peptides by cd-hit (Li & Godzik, 2006) were listed in Table S1. A total of 643,020 non-redundant peptide sequences from the three genomes (*P. patens, M. polymorpha* and *S. moellendorffii*) and 26 *de novo* transcriptomes were sorted into 31,779 orthogroups, which was reduced to 5,684 by retaining only those orthogroups that included groups representing a minimum of 10 taxa and with at least one sequence in each of the three reference genomes (*P. patens, M. polymorpha* and *S. moellendorffii*). The 5,684 orthogroups were further collapsed to retain only one representative sequence for mono- and para-phylogenetic clades that contained sequences from single taxa, and terminal branches with abnormal lengths were eliminated (Fig. S1). Finally, the 649 one-to-one orthogroups were used for species tree estimation and the remaining 5,035 multi-copy orthogroups for the identification and resolution of the phylogenetic placement of WGD events through subsequent gene tree reconstructions and reconciliation with the species tree. The masking of transcript isoforms and recent paralogs did not alter the detection of WGDs (Fig. S2 and S3) and the trimmed gene family trees removed spurious terminal branches that eliminated the inclusion of unreliable members derived from mis-assemblies (Yang & Smith, 2014).

**Fig. 1.**
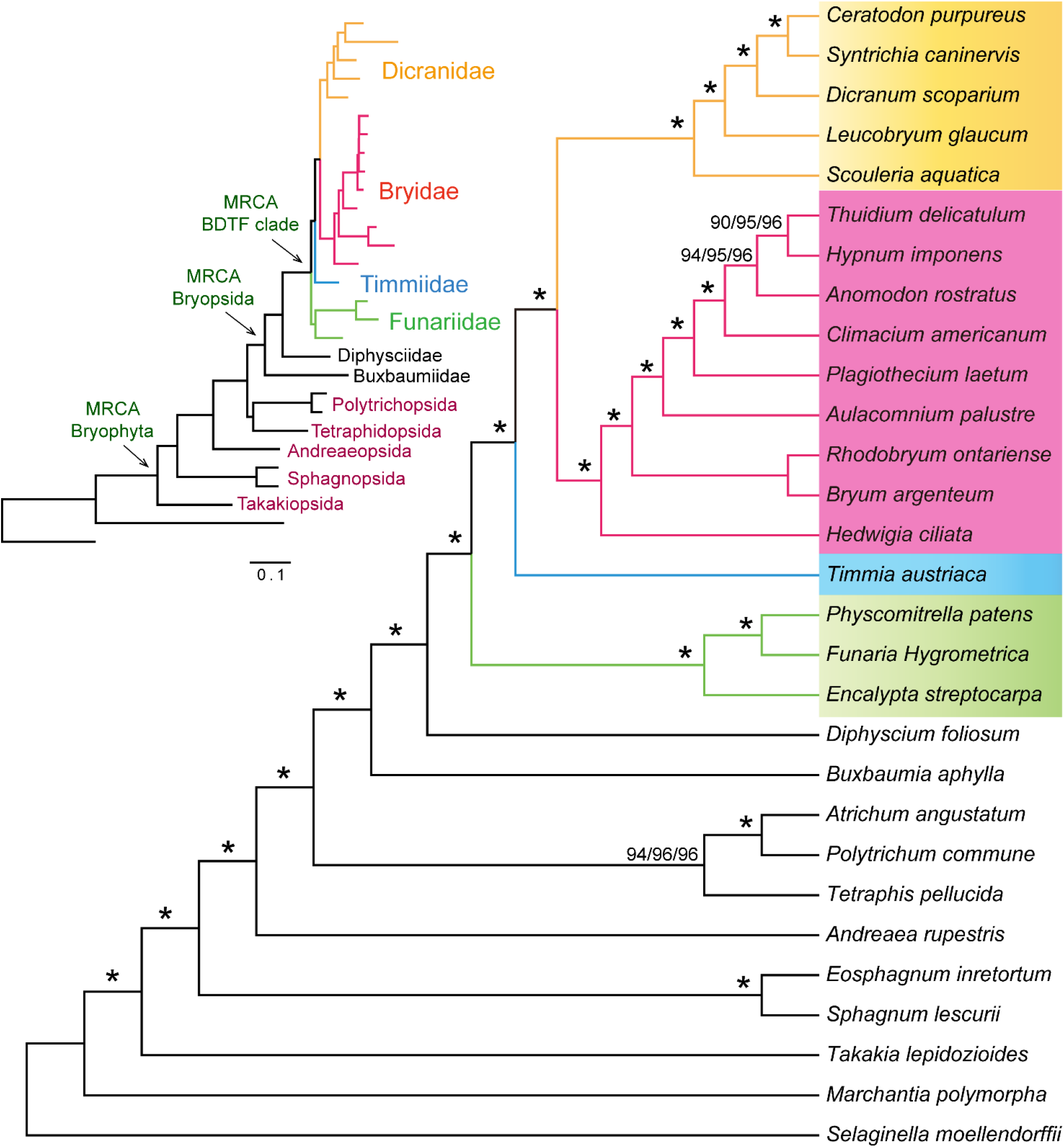
Evolutionary relationship reconstructed from concatenated alignment of the coding sequences of 649 orthogroups in 27 mosses, one liverwort and one lycophyte. Bootstrap values are labeled at the nodes and were obtained from three partition strategies: unpartitioned/(1st+2nd) and 3rd codon positions as two partitions/PartitionFinder scheme. The asterisks show nodes supported by bootstrap values of 100% (500 replicates). Branch lengths of the maximum-likelihood phylogram generated under the PartitionFinder scheme with ancestral nodes were depicted in the top left corner.

### Moss phylogeny reconstructed from single-copy orthogroups

The sequence alignments from the 649 putative single-copy gene families were concatenated to generate a sequence supermatrix that contained 833,313 sites. The contributions of each species to the single-copy orthogroups are presented in Table S2. With the exception of the placement of *Hypnum imponens* and *Anomodon rostratus* relative to *Thuidium delicatulum* within Hypnales and the clade of Polytrichopsida plus Tetraphidopsida (Fig. S4), the maximum-likelihood (ML) species tree generated by RAxML produced a well-resolved moss phylogeny with bootstrap support values (BSV) of 100% for all other nodes. Though the BSVs for these three nodes in Hypnales were not 100%, all were larger than 90% in the three partition schemes (Fig. 1). The partitioning scheme recommended by PartitionFinder produced a moss phylogeny with relatively better BSV support values for the three nodes in Hypnales nodes (BSV > 95%) supporting the phylogeny presented in Fig. 1. The Polytrichopsida plus Tetraphidopsida clade was also supported when species tree was constructed using only the 1st and 2nd codon positions in the concatenated alignment (Fig. S5), and the concatenated protein alignment (Fig. S6) also generated the identical species tree topologies with 100% BSVs.

A coalescence-based species tree was estimated from the 649 individual gene family trees using the ASTRAL-III v5.6.2 algorithm (Zhang *et al.*, 2018). With the exception of the deep detailed phylogeny in Hypnales, the ASTRAL-III species tree topologies were consistent with the phylogeny reconstructed from concatenated supermatrix (Fig. S4). The Polytrichopsida plus Tetraphidopsida monophyletic clade was supported in the ASTRAL-III analysis (BSV of 100%).

Overall, the species tree topologies were consistent with the accepted moss taxonomy (Shaw *et al.*, 2011). *Takakia lepidozioides* (Takakiopsida) were confidently placed at the basal position sister to all other mosses and followed by the Sphagnopsida. Several studies had resolved *Takakia* as sister to *Sphagnum*, e.g. (Nickrent *et al.*, 2000; Qiu *et al.*, 2006; Shaw *et al.*, 2010a), but none were supported with high confidence. Our analyses revolved Takakiopsida as a basal-most lineage of mosses with 100% bootstrap support in all ML and ASTRAL-III analyses, indicative of the high resolving power of phylotranscriptomic approach to reconstruct species phylogeny such as recently demonstrated for the phylogeny of gymnosperms (Ran *et al.*, 2018). The phylogenetic relationships for other species-poor lineages outside of Bryopsida were also well resolved in our analyses: Andreaeopsida (*Andreaea rupestris*) were basal to the Polytrichopsida plus Tetraphidopsida clade that together constituted graded sister groups to the class Bryopsida (Fig. 1).

Within the species-rich class Bryopsida (true mosses), the subclasses Funariidae and Timmiidae (*Timmia austriaca)* was recognized as graded sister groups to the clade of Dicranidae plus Bryidae. These four subclasses together constituted a large proportion (more than 95%) of modern mosses (Newton *et al.*, 2006) and demonstrated a relatively short ancestral branch length indicative of a rapid species diversification process (Fig. 1). For expediency, we refer to them as the BDTF clade (Bryidae, Dicrsanidae, Timmiidae and Funariidae). The other species-poor “basal Bryopsida” subclasses Diphysciidae (*Diphyscium foliosum*) and Buxbaumiidae (*Buxbaumia aphylla)* resolve as laddered sister groups to the BDTF clade.

The detailed deep phylogeny of the Hypnales remains a hard-resolving problem because of rapid species diversification within the group (Johnson *et al.*, 2016). Previous phylotranscriptomic studies, using similar concatenated supermatrix-based and coalescence-based strategies, also encountered inconsistent topology (Wickett *et al.*, 2014) or relatively low BSV supporting values for internal nodes (Johnson *et al.*, 2016) within the Hypnales phylogeny. The tree topology within Hypnales generated from our concatenated supermatrix was congruent to that reported in (Johnson *et al.*, 2016), but with much higher bootstrap supporting scores. The high-confidence supermatrix-based species tree topology generated here (Fig. 1) was employed in subsequent phylogenomic analyses for the detection of ancestral polyploidy events.

### Detection and phylogenetic placement of ancestral genome duplications

The *Ks* frequency plots have been used to detect signals for one or more ancestral whole genome duplications for mosses, (e.g. (Rensing *et al.*, 2007; Szovenyi *et al.*, 2015; Lang *et al.*, 2018)). Lang *et al*. detected *Ks* peak signals in most Bryopsida and Sphagnopsida species and circumscribed two rounds of WGDs based on the chromosomal syntenic structure in the *P. patens* genome (Lang *et al.*, 2018). However, detecting WGD events using paranome based *Ks* distributions is difficult (Vekemans *et al.*, 2012) and the saturation effect of *Ks* and rate heterogeneity among different gene families and lineages, among other issues, could also blur Ks-based age peaks (Jiao *et al.*, 2011). Large-scale gene tree-based phylogenomic method described in (Jiao *et al.*, 2011) avoids many of these issues and has been effectively utilized in recent ancestral genome duplication studies (Jiao *et al.*, 2012; Devos *et al.*, 2016; McKain *et al.*, 2016; Unruh *et al.*, 2018).

To utilize the benefits of the large-scale gene tree-based phylogenomic method, we examined the gene trees constructed from the 5,035 multi-copy orthogroups using the PUG algorithm (McKain *et al.*, 2016; Unruh *et al.*, 2018), based on the moss phylogeny we had constructed, to detect six focal nodes with high concentrations of gene duplications (Fig. 2). The ancestral node with the largest number of gene duplications (N20 in Fig. 2, 988 duplications with BSV ≥ 80%) was identified at the MRCA of BDTF clade. This provided evidence supporting a genome duplication event shared by all species in Bryidae, Dicranidae, Timmiidae and Funariidae, designated the ψ event or BDTF duplication hereafter. Large-scale gene duplication events were also indicated in the ancestor of the Funarioideae (N4 in Fig. 2, 794 unique duplications with BSV ≥ 80%) which was not shared with *Encalypta streptocarpa* (Encalyptaceae) and in the ancestor of Bryopsida, Polytrichopsida and Tetraphidopsida (BPT clade, N23 in Fig. 2, 265 duplications with BSV ≥ 80%). The analyses also captured the Sphagnopsida-wide genome duplication event (N1, 401 duplications with BSV ≥ 80%) and the ancestral moss-wide duplication before the diversification of all sampled mosses (N26, 357 duplications with BSV ≥ 80%) reported earlier (Devos *et al.*, 2016). Our analyses suggested that *Takakia lepidozioides* (Takakiopsida) was included in the moss-wide genome duplication event. Moreover, larger numbers of gene duplications with BSV ≥ 50% were also labeled and highlighted at each internal node, yielding consistent results with the cutoff at BSV ≥ 80% (Fig. 2). Furthermore, the large-scale gene family expansion/duplication occurring before the rapid speciation in Hypnales was also captured, however, the number of detected gene duplications are low (N14, 149 duplications with BSV ≥ 80%) and so does not strongly support a recent WGD event: more detailed analyses of gene family expansions in the Hypnales were reported in (Johnson *et al.*, 2016).

**Fig. 2.**
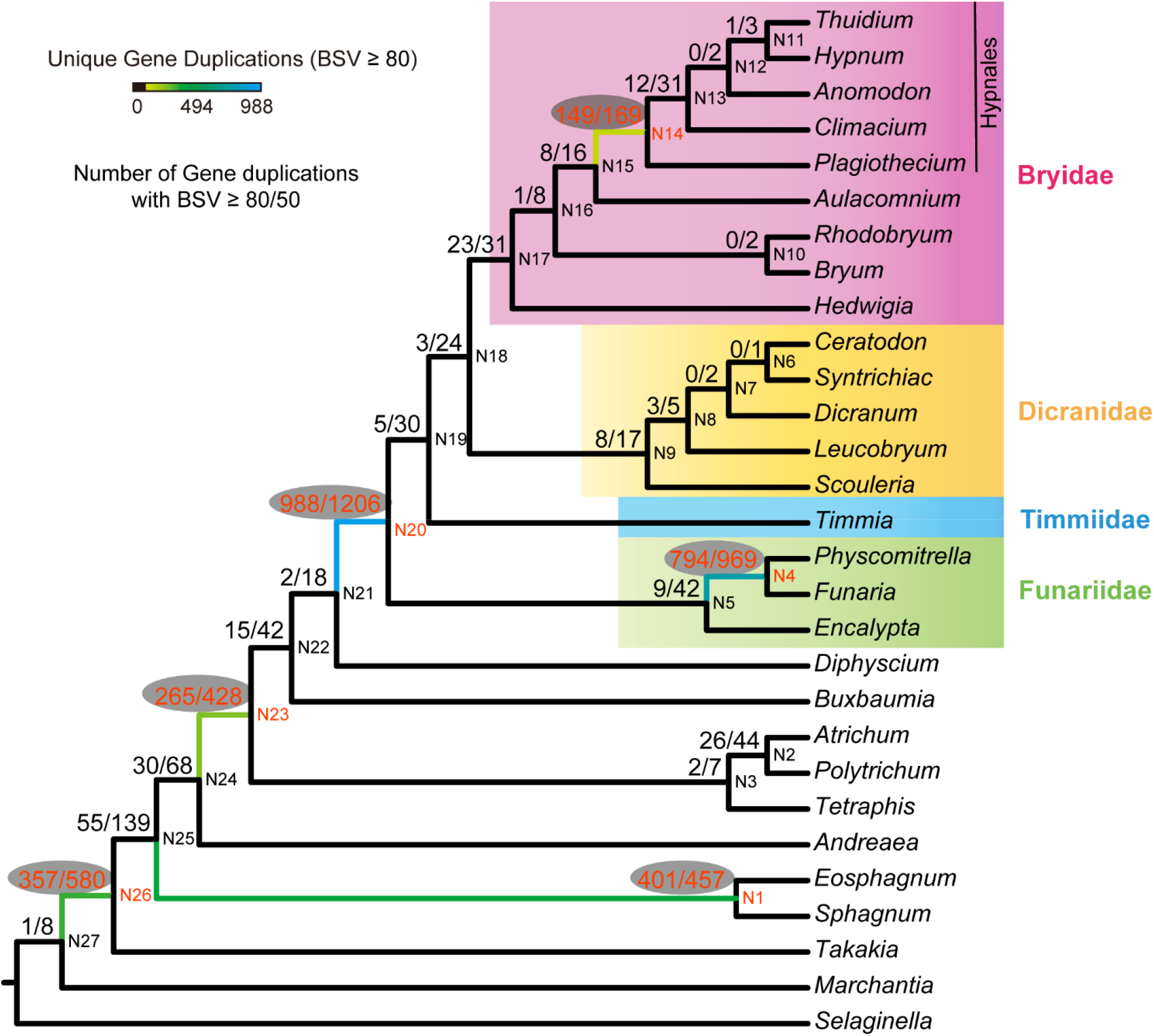
Detection of focal nodes with high concentrations of ancestral paralogous gene duplication events. Mapping results from querying paralogous pairs identified from gene-tree species-tree reconciliation using the PUG algorithm. Number of duplication nodes with bootstrap values no less than 80% and 50% were counted and labeled at corresponding ancestral nodes on the species tree. Colored branches highlighted the number of unique gene duplications as depicted in the key.

### Resolving the two WGDs in *Physcomitrella patens* utilizing phylogenetic timing of syntelogs

Phylogenetic timing of duplicated paralogs retained on genomic syntenic regions in the *Physcomitrella patens* genome were scrutinized and summarized on the moss phylogeny (Fig. 3 and Table 1). As a result, more than 95% of the analyzed syntelog pairs were phylogenetically identified to be products from the N20 (BDTF clade) and N4 (Funarioideae) duplications. Using the *P. patens* syntelogs as anchors, a total of 93 and 114 duplication nodes were detected in the ancestor to the BDTF clade with minimum BSVs of 80% and 50%, respectively. In addition, 287 and 339 duplications were circumscribed at the ancestor of Funarioideae (shared by *Physcomitrella* and *Funaria*) with minimum BSVs of 80% and 50%, respectively. While all other ancestral nodes demonstrated to contain very small number (less than 1.5%) of duplicated syntelogs (Table 1).

**Fig. 3.**
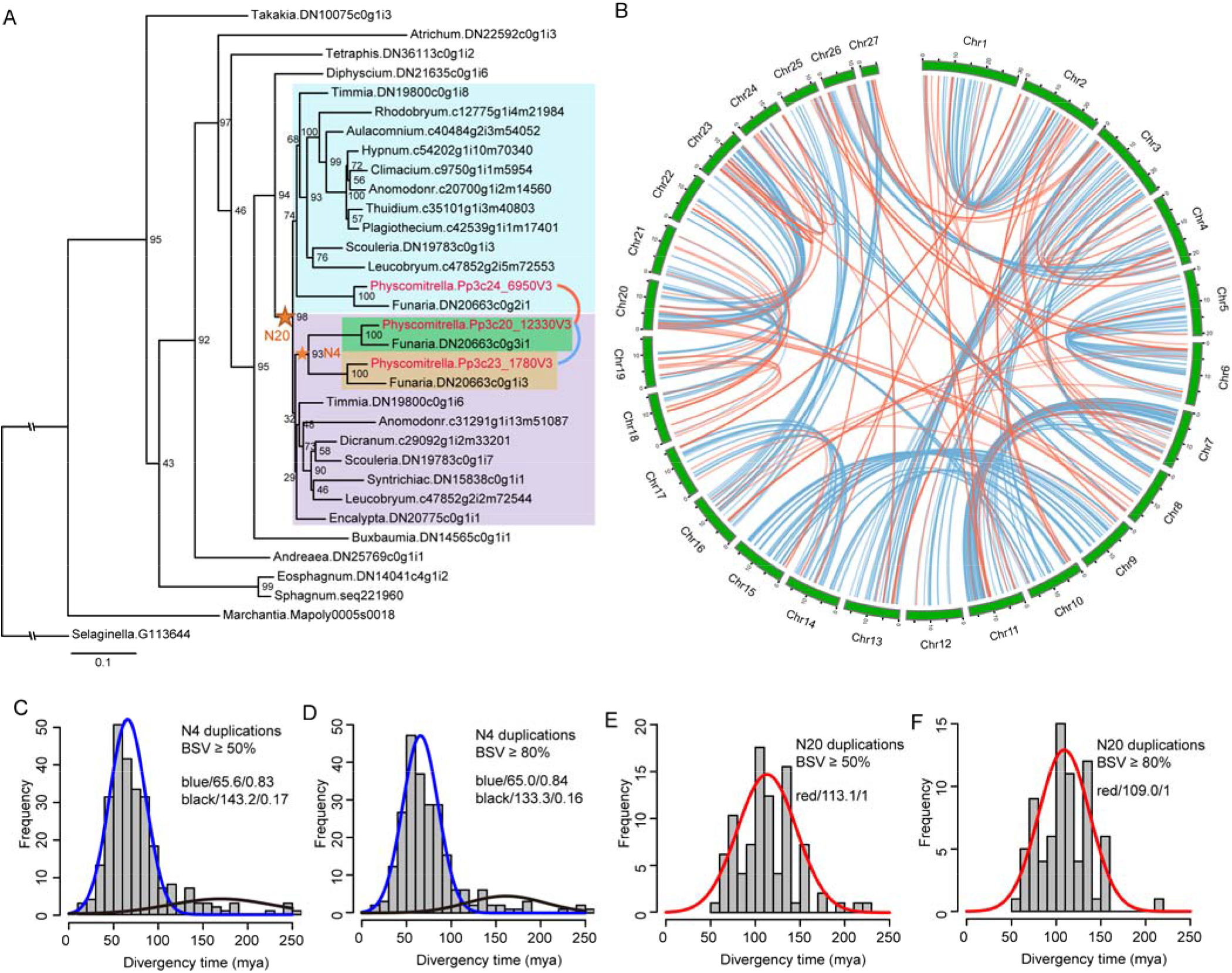
Determining the phylogenetic position and age of the two polyploidy events in *Physcomitrella patens* by tracing the phylogenetic timing of genomic syntelogs. **(a)** Exemplar maximum likelihood gene tree from OrthoGroup_1244. Two duplication nodes (with BSV ≥ 80%) were identified in this gene family phylogeny and were highlighted with stars. The paralogous pair *Pp3c20_12330V3* and *Pp3c23_1780V3* (connected by blue curve) were retained on syntenic regions in *P. patens* genome and supported the Funarioideae (N4) duplication, while the paralogous pair *Pp3c20_12330V3* and *Pp3c24_6950V3* (connected by red curve) were detected on *P. patens* genome syntenic regions and supported the ψ (N20) duplication event. **(b)** Spatial distribution for the N4 (connected by blue lines) and N20 (connected by red lines) duplicated syntelogs in the *Physcomitrella patens* genome. The phylogenetic timing for the syntelog pairs in *Physcomitrella* were inferred from gene family phylogenies (as summarized in Table 1). The Funarioideae (N4) and BDTF-wide (termed ψ event, N20) duplications were depicted as blue and red lines, respectively. **(c)** The inferred duplication times for *P. patens* syntelog pairs that support the Funarioideae-wide duplication (BSV ≥ 50%) were analyzed by EMMIX. Each statistical component was written as ‘color/mean molecular timing/proportion’ where ‘color’ is the component (curve) color and ‘proportion’ is the percentage of duplication nodes assigned to the identified component. **(d)** Age distribution and mixture model analyses of the Funarioideae duplications with BSV ≥ 80%. **(e)** Distribution of estimated duplication times from *P. patens* syntelogs that support a BDTF clade duplication (ψ event) with BSV ≥ 50%. **(f)** Distributions of inferred duplication times support BDTF clade duplication (ψ event) with BSV ≥ 80%.

**Table 1.**
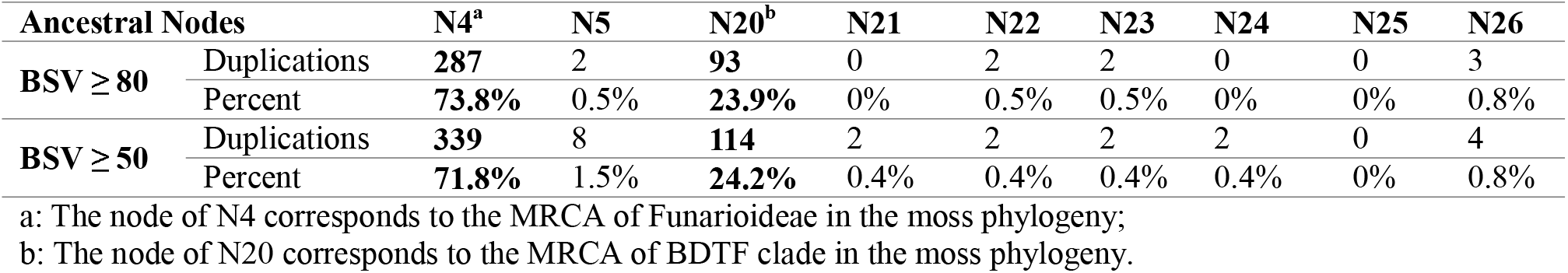
Phylogenetic timing of the *Physcomitrella patens* syntelogs inferred from gene family phylogenies

To investigate whether the phylogenetic timing of duplicated syntelogs in *Physcomitrella patens* were consistent with the two ancestral WGD events (ψ and Funarioideae duplications), we connected the syntelogs derived from N20 and N4 duplications using CIRCOS v0.69-6 (Krzywinski *et al.*, 2009) (Fig. 3a and 3b) and compared it to the *P. patens* chromosomal syntenic karyotype (Lang *et al.*, 2018). The spatial distribution of the duplicated syntelogs provided strong evidence that the older WGD1 is congruent to the BDTF clade (N20) duplication, while the younger WGD2 is consistent with the Funarioideae (N4) duplication. For example, the Chr5 vs Chr6, Chr7 vs Chr11 and Chr9 vs Chr15 are chromosome pairs representing products of the younger WGD2 events (Lang *et al.*, 2018), and they are heavily populated with connected duplicated syntelogs (blue connecting lines in Fig. 3b) that is indicative of Funarioideae (N4) duplications. Consistent phylogenetic timing evidences for the older WGD1 patterns could also be observed in the analyses, for example, the four chromosomes 18/21 and 19/22 in *P. patens* were the products derived from the two WGD events from a single ancestral chromosome, largely congruent with (((chr18,chr21)WGD2,(chr19,chr22)WGD2)WGD1) (Lang *et al.*, 2018). Duplicated syntelog pairs between Chr18 and Chr19/22 are heavily populated with red connecting lines (Fig. 3b), corresponding the BDTF clade ψ duplications at N20, consistent with the older WGD1 event. Similarly, the Chr23/20 vs Chr24, chromosomes pairs (derived from a single ancestral chromosome) duplicated from the older WGD1, are also heavily populated with syntelogs congruent with the N20 ψ duplications in gene family phylogenies.

The phylogenomic analyses above provided robust phylogenomic evidence that the ancestral ψ genome duplication event occurred prior to the diversification within the BDTF clade and the data is consistent with the reported WGD1 event in *Physcomitrella patens* (Lang *et al.*, 2018) and the Pp-WGD2 event was circumscribed in the ancestor of the Funarioideae.

### Molecular dating of the Funarioideae and ψ duplications from *Physcomitrella patens* syntelogs

Single gene family trees may contain more than one duplication nodes (e. g. Fig. S3 and Fig. 3a), so we counted and estimated the ages of each duplication node to estimate the ages of the Funarioideae and ψ duplication events. Gene trees that did not pass the cross-validation and gradient check procedure in r8s (Sanderson, 2003) were discarded. Finally, divergency times were estimated for 268 out of the 339 duplication nodes with a minimum BSV of 50% for the Funarioideae duplication, and 89 of the 114 duplication nodes in agreement with the BDTF clade duplication that were supported by a minimum BSV of 50%. The frequency distribution of divergency times of the *P. patens* syntelog pairs that traced the Funarioideae-wide duplications exhibited a peak at 65.8 ± 2.6 mya (95% confidence interval) (Fig. 3c), and the distribution of ψ duplication ages exhibited a peak at 113.1 ± 6.9 mya (Fig. 3e). The age frequency distributions for ancestral duplications with higher BSV thresholds at 80% were also evaluated and a total of 229 and 79 duplication age estimates were obtained for the Funarioideae and ψ duplications, respectively. The estimated ages of the Funarioideae duplications exhibited a significant primary component around 65.6 ± 2.7 mya (Fig. 3d), and a significant age component around 109 ± 6.6 was detected for the ψ duplications (Fig. 3f).

In both analyses, using BSV thresholds at 50% and 80%, the age distributions of the ψ duplications were modeled to a single peak that exhibits relatively higher variation, whereas conspicuous peaks circa 65 mya were observed for the Funarioideae duplications. However, the mixture model implemented in EMMIX (McLachlan *et al.*, 1999) also detected older component with a very low proportion in the Funarioideae duplication ages. The tree uncertainty, incomplete lineage in some of the gene trees and some extreme evolutionary rate heterogeneities in gene families (drastically distinctive habitats from desert to bogs) may result in a small proportion of age outliers the age estimations. We cannot either exclude the possibility of the inclusion of some older segmental or small-scale duplications (SSDs). Whereas the major conspicuous component in the age frequency distribution around the Cretaceous-Paleogene (K-Pg) boundary was well-circumscribed in the analyses and consistent with the previous study (Vanneste *et al.*, 2014).

To evaluate the sequence divergency of the syntelogs following paleopolyploidy, we generated and analyzed the *Ks* frequency distributions for the syntelog pairs that traced the Funarioideae and ψ duplication events in the gene family phylogeny (Fig. 4). The observed *Ks* frequency distribution peaks were consistent with the molecular dating results as described above, the Funarioideae and ψ duplications are in parallel with major *Ks* peaks around 0.6 (Fig. 4a and 4b) and 0.9 (Fig. 4c and 4d), respectively.

The synonymous divergency curves for syntelog pairs derived from both Funarioideae and ψ duplications were obviously right-skewed and the *Ks* frequency distributions could be modeled with two major components containing one “younger” peak component and one “older” right-skewed component (blue and green for Funarioideae duplications in Fig. 4a and 4b, red and purple for ψ duplications in Fig. 4c and 4d). The analyses using the BSV threshold at 50% for each component was described by “color/mean/proportion”, in agreement with previous analyses (Jiao *et al.*, 2011; Jiao *et al.*, 2012). The *Ks* frequency distribution of syntelog pairs derived from the Funarioideae duplications were modeled with three components: blue/0.59/0.67, green/0.88/0.27 and black/1.36/0.06 (Fig. 4a) and the *Ks* frequency of ψ event syntelogs contained two statistically significant components: red/0.9/0.775 and purple/1.5/0.225 (Fig. 4c). Thus, the “green” component (Fig. 4a and 4b) in the Funarioideae duplications could severely blur the major component of the ψ duplications as shown in Fig. 4e. This synonymous distance distribution pattern would make it difficult to infer ancestral genome duplications directly from the *Ks* distribution without the phylogenetic timing information. We also analyzed the *Ks* frequency distribution for all syntelogs in *P. patens*, the four major *Ks* components were recaptured (Fig. 4f) and were similar to that reported in (Lang *et al.*, 2018).

We described each component using “component no./mean/proportion”: 1/0.56/0.21, 2/0.76/0.33, 3/1.09/0.41 and 4/1.8/0.05, and according to the *Ks* distribution analyzed above, the youngest component (1/0.56/0.21) is largely in parallel to the Funarioideae duplication event. However, the other two major components (2/0.76/0.33 and 3/1.09/0.41) could be a mixture of both Funarioideae and ψ duplications.

**Fig. 4.**
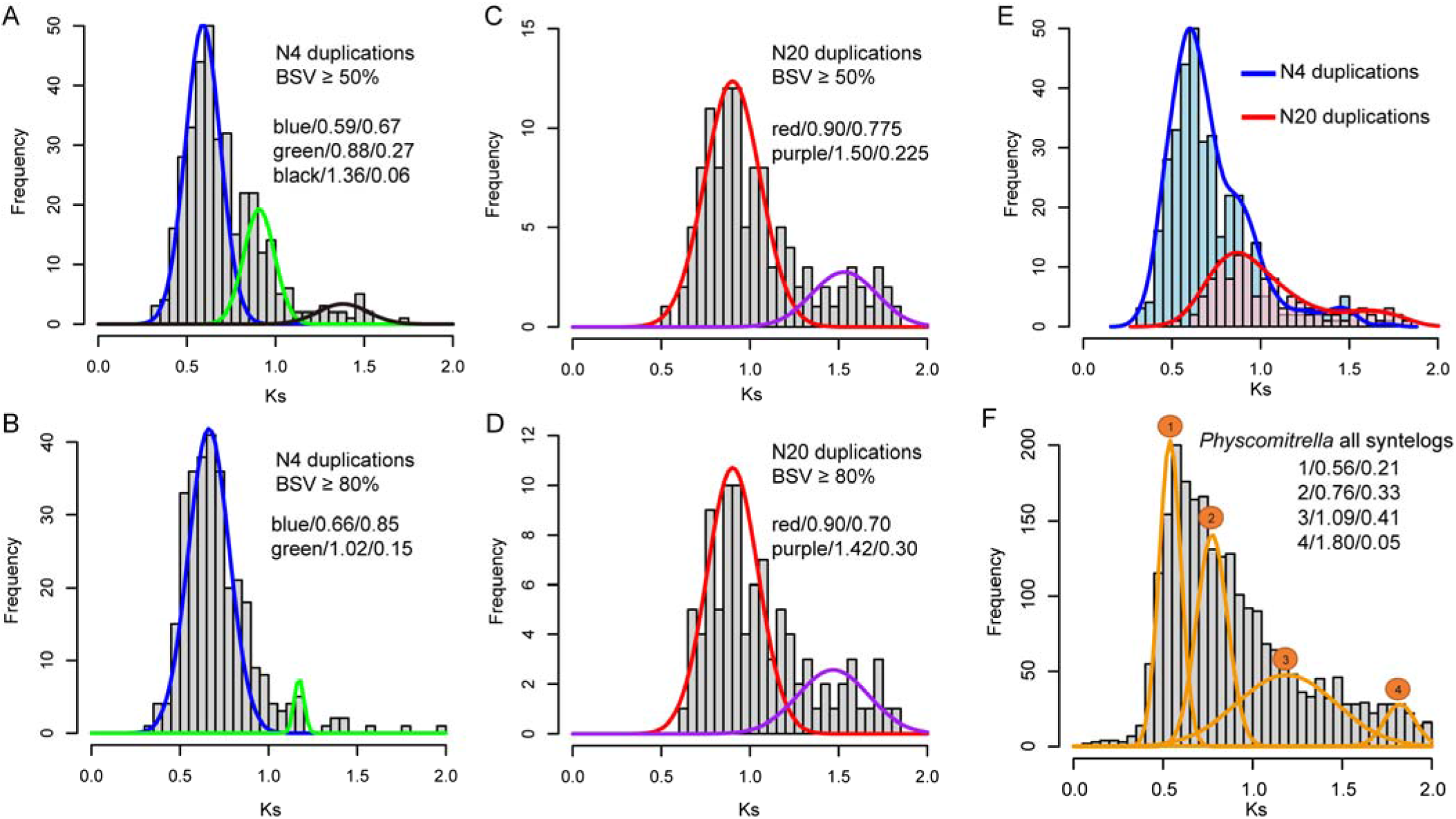
*Ks* distribution analyses of *Physcomitrella patens* syntelogs. Pairwise synonymous distances for duplicated paralogs retained on syntenic blocks in the *P. patens* genome were analyzed in combination with their phylogenetic timing information. **(a)** *Ks* distribution from 339 *Physcomitrella* syntelog pairs supporting a Funarioideae duplication (BSV ≥ 50%) in the phylogenetic analysis. **(b)** *Ks* distribution of 114 *Physcomitrella* syntelog pairs supporting a BDTF clade duplication (BSV ≥ 50%) in the phylogenetic analysis. **(c-d)** *Ks* distributions of *Physcomitrella* syntelogs supporting a Funarioideae (287 pairs) and BDTF clade (93 pairs) duplications in the phylogenetic analyses with BS ≥80%. **(e)** Contrasting plot for the *Ks* distributions of *Physcmitrella* syntelogs supporting Funarioideae (blue) and BDTF clade (red) duplications (BSV ≥ 50%). **(f)** *Ks* distribution of all paralogous pairs identified from *Physcomitrella patens* genome syntenic block analysis.

### Nature of the ψ duplication event

We have dated the age distribution using the *P. patens* ψ-derived syntelogs to be around 100-110 mya, however with only a small number of aged duplication nodes (89 and 79 with BSV cutoffs at 50% and 80%, respectively), it is possible this may add an additional source of error and could have affected the subsequent mixture modeling analyses (Morrison, 2008; Devos *et al.*, 2016). To overcome the small sample size and to further characterize the ψ event, we dated all 1,206 (BSV ≥ 50%) and 988 (BSV ≥ 80%) duplication nodes that traced the ψ event (Fig. 2), and we obtained age estimations for 787 and 935 duplication nodes respectively using the r8s package (Sanderson, 2003).

The derived age distributions from the expanded analysis were similar to the age distributions obtained from *P. patens* syntelogs with the majority of the derived divergency ages clustering around ~100 mya (Fig. 5a and 5b). However, the divergency ages for the ψ duplication event contained two significant major components consistent with the syntelog *Ks* frequency distribution patterns. For clarity, we formulated each significant component in the age distribution as “color/95% CI of the mean age/proportion”. The age frequency distribution for ψ duplications (BSV ≥ 50%) contained three significant components: red/86.7 ± 1.2/0.56, purple/136.8 ± 2.4/0.36 and black/216.5 ± 15.2/0.08 (Fig. 5a). Similarly, three significant components were also identified for ψ duplications with BSV ≥ 80%: red/86.7 ± 1.3/0.617, purple/136 ± 2.4/0.345 and black/226.7 ± 16.9/0.038 (Fig. 5b). The patterns of the duplication age distribution in this analysis suggests that about 30% of the ψ gene duplications occurred around 136 mya with the majority (~60%) estimated to be duplicated around 87 mya.

**Fig. 5.**
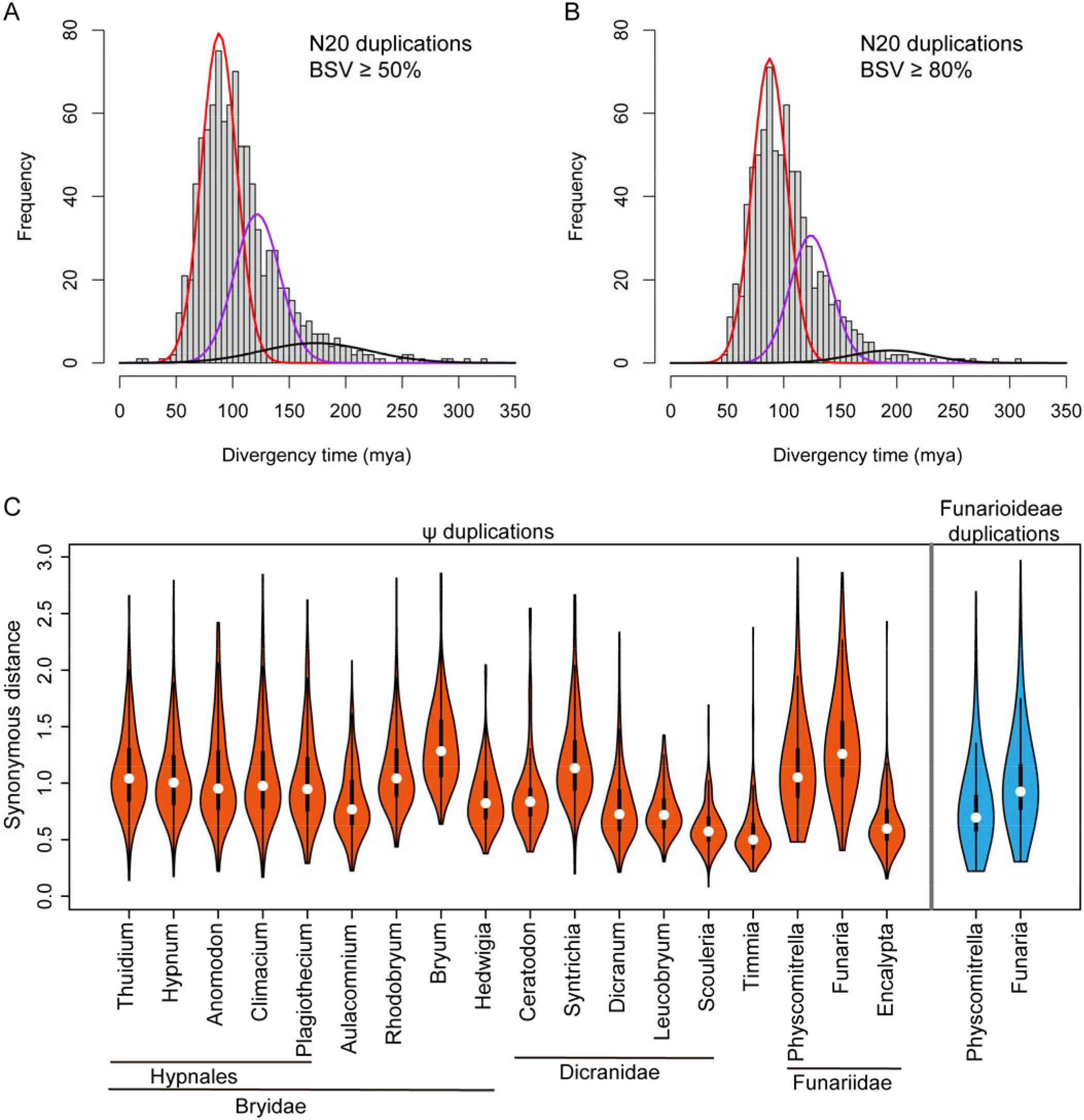
Characterization of the ψ event duplications in the common ancestor of BDTF clade. All estimated paralogs constituting the N20 duplications (ψ event) were included in the age EMMIX analyses. Each significant component is depicted as “color/95% CI of the mean age/proportion” where “color” is the component (curve) color and “proportion” was the estimate of mixing proportion for each component. **(a)** Age distribution of all N20 duplications with BSV ≥ 50% indicates three significant components: red/86.7 ± 1.2/0.56, purple/136.8 ± 2.4/0.36 and black/216.5 ± 15.2/0.08. **(b)** Age distribution of all N20 duplications with BSV ≥ 80% indicates three significant components: red/86.7 ± 1.3/0.617, purple/136 ± 2.4/0.345 and black/226.7 ± 16.9/0.038. **(c)** Violin plot contrasting *Ks* distributions of the ψ and Funariideae duplicated paralogs in different mosses indicated nucleotide substitutional rate variations. All inferred paralogous pairs tracing the ancestral ψ and Funarioideae genome duplications (BSV ≥ 50%) were analyzed to calculate nucleotide substitutional rates among BDTF clade mosses.

To further validate the “two-component” divergency pattern of the ψ genes, we extracted all paralogous pairs with coalescence node at N20 (BSV ≥ 50%) that tracing the ψ events in gene family trees. *Ks* values were estimated for these “pure” sets of ψ duplicated paralogous pairs which are largely free of background duplications. In parallel with the age estimations (Fig. S7), *Ks* frequency distributions for almost all BDTF clade species were modeled with two significant components (Fig. S7, A-R). For example, consistent to the dating analyses based on *P. patens* syntelogs (Fig. 4c and 4d), the two major components ‘red/0.9/0.51’ and ‘purple/1.26/0.4’ were also identified in the *Ks* distribution for the ψ duplicated genes inferred from gene trees (Fig. S7L). While for all the inferred *P. patens* paralogous pairs with coalescence node at N4 (BSV ≥ 50%) tracing the Funarioideae duplication event, two major components ‘blue/0.63/0.66’ and ‘green/0.99/0.3’ were identified (Fig. S7T), which was also consistent with the syntelog-based analyses described above. The intriguing pattern in the *Ks* frequency distribution may suggested that a small proportion of moss genes has accumulated more synonymous mutations.

However, the heterogeneity in evolutionary rate between different gene families may not fully explain the distinctive “two components” in the age distribution of the ψ derived duplicates. Other events may have contributed to the ‘older’ component (~136 mya). The two components are not the result of two recurrent genome duplication events or a triplication event as that would have been uncovered in the genomic synteny depth analysis for *P. patens* (Lang *et al.*, 2018). It is possible that segments of the genome were duplicated prior to the merging of the subgenomes that constitute the ψ polyploidy event and would thus contribute to the older component. However, this is unlikely as we did not observe specific genomic segments that were significantly older than others. It is more likely that the ψ duplication was an allopolyploidy event resulting from the merging of diverged dioecious female and male gametophyte genomes, as suggested by Rensing *et al*. (Rensing *et al.*, 2007), within which the divergencies were preexisting before the WGD event. This then indicates that the dated nodal ages traced the divergency time of the separation of the two sub-genomes and not the polyploidy event itself (Doyle & Egan, 2010). We favor this hypothesis because in this scenario the allopolyploidization of dioecious gametophytes could yield a monoecious plant which could enhance the dispersal of breeding populations (Rensing *et al.*, 2007), and thus provide extra ecological advantages for polyploids.

Although we support the ψ allopolyploidy hypothesis, we cannot fully rule out the possibility that the older component of the ψ duplications might be the result of errors introduced in the large-scale molecular dating procedure. Although the detection of the major age components constituting the duplication events is clear in the genome-wide analyses for thousands of gene families we performed, the setting of the ancestral fixed age constraints at the roots of the gene trees and the possibility that tree topologies and branches length estimations may introduce some abnormal age estimations.

It has been proposed polyploidy in mosses could be frequent and independent events in different species and isolated individuals (Rensing *et al.*, 2007), but even if some mosses escaped the ψ and/or Funarioideae events, such cases would be very rare, and the evolutionary significance of the ψ event remains valid. The sequencing of other moss genomes (specifically a species in the BDTF clade) will improve the inter-genome synteny comparisons and will clarify the timing and nature of the ψ event since at present the conclusion is that the ψ duplication in *P. patens* was superseded by the more recent Funarioideae WGD event. The generation of more sequence data in moss species that represent closer sister relationships to the BDTF clade, e.g. species in Gigaspermaceae (Medina *et al.*, 2018), may provide a more precise phylogenetic position for the ψ event.

### Evolutionary rate variations among mosses

Variations in nucleotide substitutional rates in the evolution of angiosperm genomes is common and have been linked to distinctive life histories (Smith & Donoghue, 2008), but this has been rarely recognized and studied in the moss lineages. We noticed that there were distinctive *Ks* peaks for the shared ψ event that ranged from 0.45 (*Timmia austriaca)* to 1.32 (*Bryum argenteum*) (Fig. S7, A-R), suggesting that there are substantial evolutionary rate variations among the different lineages following the ψ event. Setting the ψ event shared by all BDTF clade species as the ancestral calibration point, we compared the nucleotide substitutional rates among the different lineages using *Ks* estimations of ψ paralogs (Fig. 5c).

We did not observe conspicuous lineage-wide nucleotide evolutionary rate shifts by inspecting the *Ks* distributions of the ψ paralogs, but there were significant within-lineage differences in evolutionary rates (Fig. 5c). The species within the Hypnales clade, within the Bryidea, exhibited similar substitutional rates of the ψ event paralogs, but for *Bryum argenteum* the rate was significantly elevated. A significant elevation of the evolutionary rate was also observed in the Dycranidae (*Syntrichia caninervis*), and in the Funariideae (*Funaria hygrometrica)*. Each of the three species, *Bryum argenteum, Syntrichia caninervis, and Funaria hygrometrica*, exhibited major *Ks* peaks (red components in Fig. S7) that exceeded 1.0 *Ks* for the ψ event. In contrast, significantly slower nucleotide substitutional rates could be observed for *Timmia austriaca, Encalypta streptocarpa* and *Scouleria aquatica*, all of which exhibited major *Ks* peaks (red components in Fig. S7) smaller than 0.6.

### Two old large-scale duplication events in mosses

Devos *et al*. reported an ancestral moss-wide polyploidy event (more than 300 mya) that was shared by the Sphagnopsida and *P. patens* (Devos *et al.*, 2016), but the precise age of this paleopolyploidy event was not determined. Two ancestral large-scale duplications (at nodes N23 and N26 in Fig. 2), predating the diversification of Bryopsida, were conspicuous in our analysis. To estimate the ages for these two large-scale duplication events, we analyzed the frequency distribution of the ages for the duplication nodes as described in (Jiao *et al.*, 2011).

Age estimations for 304 (BSV ≥ 50%, 193 with BSV ≥ 80%) duplication nodes tracing the duplication event before the diversification of Polytrichopsida + Tetraphidopsida and Bryopsida (the BPT clade) but after the separation of *Andreaea* were obtained and the duplication ages were well clustered, with a conspicuous peak, around 150-200 mya (Fig. 6a and 6d). Similarly, frequency distributions from the estimated ages tracing the moss-wide duplications, for 448 (BSV ≥ 50%,) and 281 (BSV ≥ 80%) duplication nodes, rendered modal peaks around 300-350 mya at both 50% and 80% BSV cutoffs (Fig. 6b and 6e). The combined age frequencies obtained from the two duplication events analyzed using EMMIX rendered two modal peaks at around 180 mya and 320 mya (Fig. 6c and 6f). The formulas mentioned above were used to describe the bimodal age duplications with BSV ≥ 50%: yellow/181.9 ± 4.9/0.4 and brown/329.4 ± 3.6/0.6 (Fig. 6c) and with BSV ≥ 80%: yellow/180.4 ± 5.3/0.43 and brown/316.5 ± 5.3/0.57 (Fig. 6f).

**Fig. 6.**
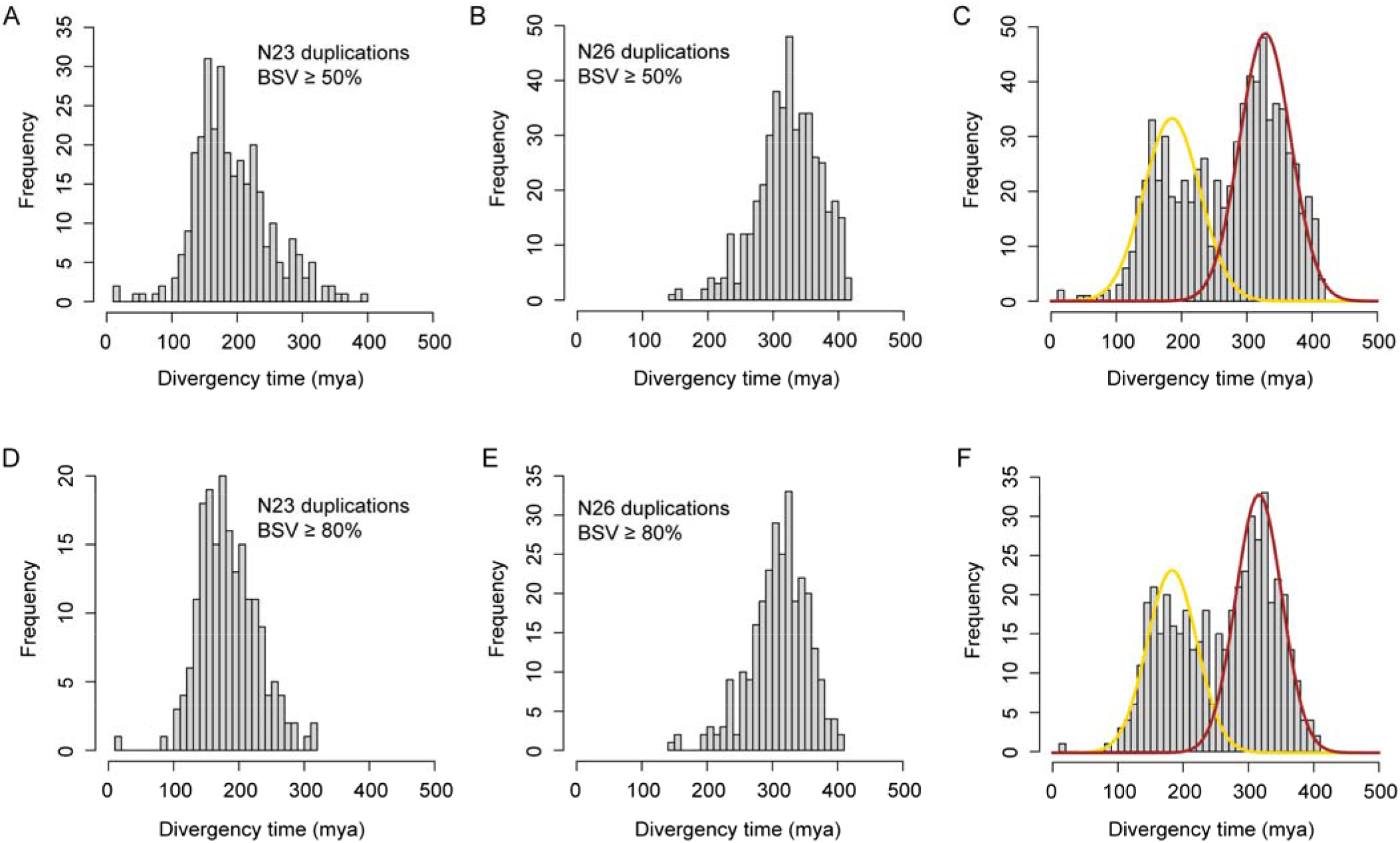
Age distribution of the two ancestral large-scale duplication events shared by all bryopsida species. **(a)** Age distribution of the ancestral duplications shared by Bryopsida, Polytrichopsida and Tetraphidopsida (BPT clade, N23) with BSV ≥ 50%. **(b)** Divergency age distribution of the ancestral moss-wide duplications (N26) with BSV ≥ 50%. **(c)** Inferred divergence times for the ancestral duplication nodes (BSV ≥ 50%) were analyzed by EMMIX and each component was described as “color/95% CI of the mean age/proportion” where “color” is the component (curve) color and “proportion” was the estimated mixing proportion for each component, two significant components were identified: yellow/181.9 ± 4.9/0.4 and brown/329.4 ± 3.6/0.6. **(d-e)** Corresponding divergency time distributions of the ancestral duplication nodes with BSV ≥ 80%, and **(f)** two significant components were identified: yellow/180.4 ± 5.3/0.43 and brown/316.5 ± 5.3/0.57.

The finite mixture model analyses confirmed the two ancestral duplications in mosses: one moss-wide duplication that occurred around 329 mya and a duplication that occurred in the ancestor of BPT clade that had an estimated age of 182 mya. The estimated age for the moss-wide duplication is largely congruent with previous reports(Shaw *et al.*, 2010b; Devos *et al.*, 2016; Morris *et al.*, 2018) and the estimated age for the moss-wide duplication event is tightly associated with the fossil record of Sphagnales at around 330 mya (Neuburg, 1960; Ignatov, 1990), though this fossil calibration was not employed in our molecular dating procedure.

Duplicated genes originated from the ψ and Funarioideae events remain pervasive in the *P. patens* genome (Lang *et al.*, 2018), however, the paralogous pairs that survived the much older duplication events would be rarer. In our dataset, we observed paralogous pairs in several transcription factor families that survived the moss-wide and BPT duplication events.

For example, the genes clustered in orthogroup_243 encode bZIP transcription factors, and *P. patens* genes were found in both of the child clades that constituted a moss-wide duplication. The more recent duplications (e.g. the ψ and Funarioideae events) that further augmented the family members were also observed in the gene family phylogeny (Fig. S8). Orthogroup_362 encodes the Nin-like transcription factors where moss-wide and BPT duplication nodes were well-supported by the tree phylogeny (Fig. S9). Similarly, the WRKY (Fig. S10) and SBP (Fig. S11) transcription factors in *P. patens* also survived the ancestral moss-wide duplication event. Besides transcription factors, the genes in orthogroup_462 encodes homologs of VAD1 (Vascular Associated Death1), a regulator of cell death and defence responses in vascular tissues (Lorrain *et al.*, 2004), also retained duplicated genes derived from the moss-wide and BPT duplications (Fig. S12). In all cases (Figs. S8-S12), the more recent ψ and Funarioideae duplication events contributed extra family members for specific clades in the gene family phylogeny. The preservation of the paralogs derived from ancient duplication events, and the recurrent acquisition of family members exert evolutionary and ecological significance for functional renovations and the species radiation process.

### Revisiting the time frame for BDTF clade diversification

The time frame for the diversification of BDTF clade (congruent with the separation of Funariidae and Bryidae/Dicranidae) is controversial; ranging from more than 268 mya based on an isolated fossil evidence calibration (Newton *et al.*, 2006; Morris *et al.*, 2018) to around 120 mya (Larsén & Rydin, 2016). Other studies have suggested a timing of around 190 mya (Fiz-Palacios *et al.*, 2011; Magallon *et al.*, 2013) and more recently a phylogenomic study in Funariaceae suggested a species diversification event around 95-120 mya that resulted in the separation of Timmiidae and Funariidae of the BDTF clade (Medina *et al.*, 2018). In this study we generated robust phylogenomic evidence for the ψ paleopolyploidy event, that affected the four Bryopsida subclasses Bryidae, Dicranidae, Timmiidae and Funariidae (BDTF clade) and that occurred after the separation of the species-poor subclasses of Buxbaumiidae and Diphysciidae. Based on more than 5,000 gene family phylogenies and the molecular dating of more than 1,000 duplication nodes, the age of the polyploidy event was circumscribed to be around 87 mya, which predates the diversification of the BDTF clade.

We noticed that some fossil records with much older ages that were identified as pleurocarpous mosses (Bell *et al.*, 2007; Christiano De Souza *et al.*, 2012; Shelton *et al.*, 2015), but ‘pleurocarpous characters’ were probably not a definitive character and we cannot exclude the possibility that these morphological characteristics has evolved in these ‘old world’ moss lineages, which phylogenetically represented basal groups instead of within the crown group of BDTF clade.

Intriguingly, a moss fossil with an age around 330 mya were assigned to Sphagnales (Neuburg, 1960; Ignatov, 1990; Morris *et al.*, 2018), however, ancient large-scale duplication events in the ancestor of Sphagnopsida were unveiled using similar phylotranscriptomic strategy with estimated ages around 197 millions years (or alternatively two episodes aound 218 mya and 112 mya) (Devos *et al.*, 2016), which are much younger than the fossil.

These evidences suggest primitive moss linages belonging to the ‘old world’ may have evolved some morphological characteristics that resemble modern mosses, but does not necessarily exclude the possibility of a relatively younger origin (or common ancestor) of modern mosses.

To further explore the age of the BDTF clade diversification, we calculated the frequency distribution for the orthologous synonymous divergency (Ks) between Bryidae/Dicranidae/Timmiidae and *Physcomitrella*, and used modal *Ks* values as proxies, as described previously(Blanc *et al.*, 2003), for the age estimation of speciation (Fig. S13). Consistent with the *Ks* peak around 0.9 for the ψ event in *P. patens*, modal *Ks* values corresponding to species separation of BDTF clade ranged from 0.543 (*Tammia* vs *Physcomitrella)* to 0.864 (*Bryum* vs *Physcomitrella*), suggesting that the diversification of BDTF clade was much younger than previous estimations that were based on a limited number of marker genes. We further generated an ultrametric species tree using the best maximum-likelihood tree generated from concatenated alignments and this indicated a diversification age for the BDTF clade of around 80 mya (Fig. 7), which is consistent with the 87 million-years-old ψ event. This result is also in agreement with a recent study that concluded that most extant mosses were products of post-Mesozoic diversification bursts (Laenen *et al.*, 2014).

**Fig. 7.**
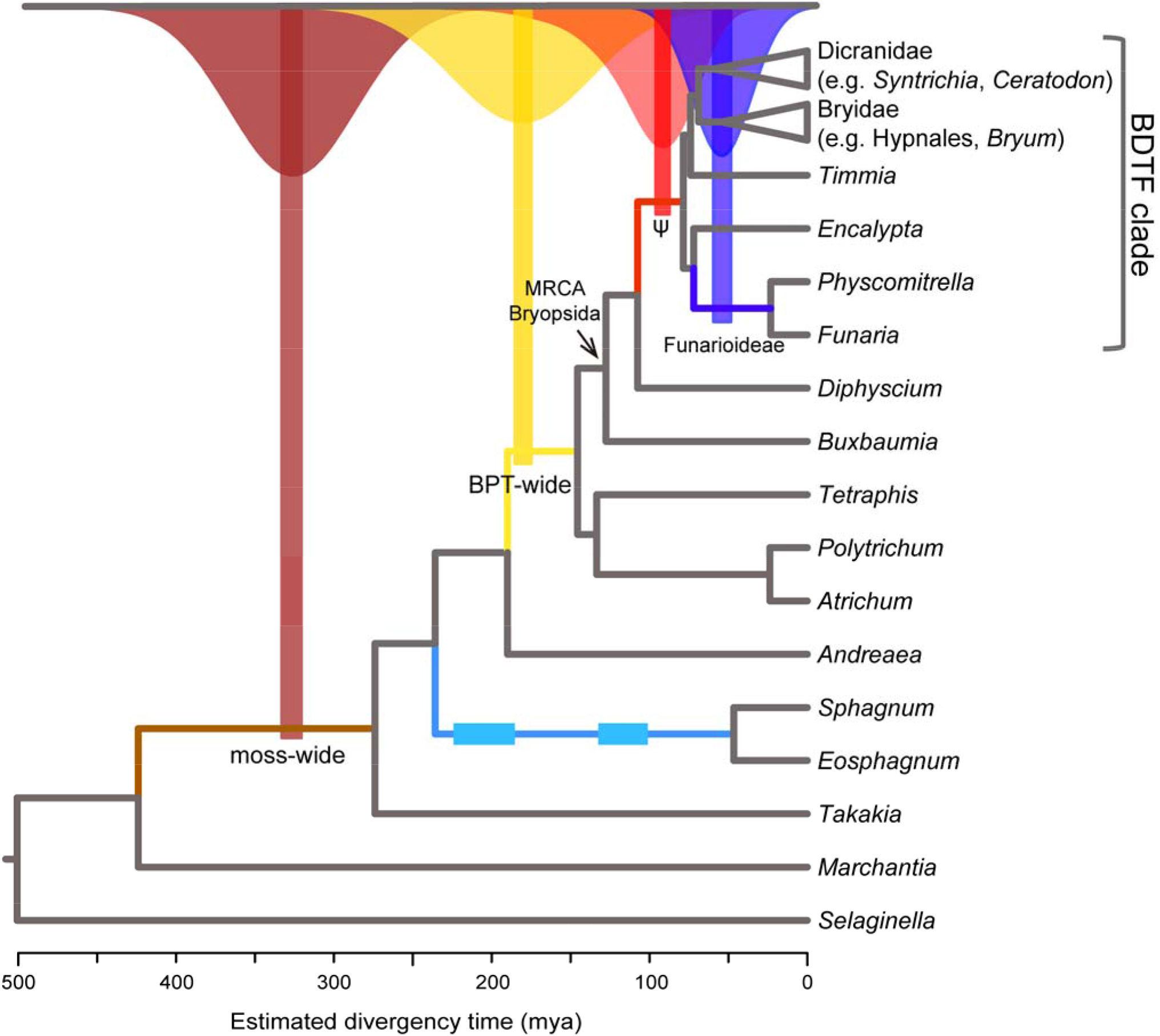
Ancestral polyploidy events in mosses. Four paleopolyploidy events in mosses were elucidated by phylogenomic and molecular dating analyses, each duplication event was represented by colored duplication age components and mapped on the collapsed species tree. The ages of nodes were estimated using r8s from the maximum-likelihood species tree constructed from concatenated alignments. The ψ allopolyploidy event was represented by its kernel density plot for gene duplication ages; while the other three polyploidy events were represented by respective Gaussian components from their respective age distributions. The polyploidy events in peatmosses were represented by two rectangles, but the actual number (one or two) of polyploidy events remains unclear.

### Implications of the ancestral large-scale duplications in mosses

Paleopolyploidy events provide substantial raw genetic materials for evolution and are recognized as important driving forces in species radiation (Schranz *et al.*, 2012; Vanneste *et al.*, 2014; Van de Peer *et al.*, 2017), though it was suggested that polyploidy could be an evolutionary ‘dead end’ (Mayrose *et al.*, 2011; Arrigo & Barker, 2012). The ψ event described here represents an example of what the WGD-RLT (Radiation Lag-Time) model has generated (Schranz *et al.*, 2012). The data we have generated indicates that the ψ event provided the impetus for a major species radiation in the Bryopsida. This is strengthened by the observation that the early-diverging Bryopdisa lineages (Diphysciidae and Buxbaumiidae) that are demonstrated to be outside of the ψ event are species-poor, and the BDTF clade, containing more than 95% of current moss species (Newton *et al.*, 2006) experienced the ψ event. Similar examples are also circumscribed by the GAMMA hexapolyploidy event in core-eudicots (Jiao *et al.*, 2012; Vekemans *et al.*, 2012).

However, unlike the GAMMA event in the core-eudicots where triplicated syntenic regions were found in the *Vitis* genome (Jaillon *et al.*, 2007), the ψ event in the mosses only generated duplicated syntenic blocks (as seen in the *P. patens* genome). The ψ event in the *P. patens* lineage likely resulted in a duplication of the seven ancestral moss chromosomes into the current 13 (including one chromosome loss) intermediate chromosomes (Lang *et al.*, 2018).

Our phylogenomic analyses circumscribed the *Physcomitrella patens* WGD2 as a Funarioideae duplication event that was shared by both *Physcomitrella patens* and *Funaria hygrometrica*. However, some *F. hygrometrica* accessions have been reported to contain 14 or 56 chromosomes and 51% of the accessions contained 28 chromosomes (Fritsch, 1991). The *P. patens* accession (Gransden 2004) has 27 chromosomes (Lang *et al.*, 2018) and some isolates of *P. patens* have 14 or 28 chromosomes (Engel, 1968), and polyploidization were frequently observed during transfection of *P. patens* protoplasts with exogenous DNA (Schween *et al.*, 2005). This suggests that some species with the Funarioideae may have escaped this duplication event or that independent polyploidization events in individuals, individual species or genera are common in the mosses.

The observations mentioned above (though focused on the Funarioideae event) also make it possible that some BDTF clade species might have escaped the ψ event, which still need to be tested. This is especially true when the transcriptomic/genomic data becomes available for *F. hygrometrica* accessions with low (e.g. n = 7) chromosome numbers, though low chromosome numbers alone could be unequivocal evidence for escaping the ancestral WGD, e.g. the five *Arabidopsis thaliana* chromosomes contained three WGDs, whereas the ancestral eudicot genome was reconstructed as seven chromosomes(Salse *et al.*, 2009). In this study we have included representative species from all the BDTF subclasses in Bryopsida, and none of them had escaped the ψ event, which is further augmented and supported by paranome-based *Ks* frequency distribution for more mosses supplemented in (Lang *et al.*, 2018).

Although the polyploidy events reported for the seed plants lineages (termed ξ, ~319 mya) and separately in the angiosperms (termed ε, ~192 mya) (Jiao *et al.*, 2011) are somewhat controversial(Ruprecht *et al.*, 2017), the ancestral large-scale gene duplications in mosses (moss-wide/329 mya, BPT-wide/182 mya and ψ/87mya) appear to be robust (Fig. 7). Our data tends to in agreement with the conclusions generated by Jiao *et al*. (Jiao *et al.*, 2011) in that the duplication events in the mosses either just predate or are just subsequent to the reported events in seed plants and are very close in geological time. This does not appear to be a consequence of the fact that our phylogenomic analyses were focused only on mosses and only the tracheophyte *Selaginella moellendorffii* was included as outgroup. The apparent co-occurrence of ancestral polyploidy and species diversification for the land plants may suggest possible ecological linkages that have orchestrated the evolution of both seed plants and mosses.

## Conclusions

A robust moss phylogeny was constructed using an extensive matrix of putative one-to-one orthogroups derived from both existing and derived transcriptomic data. Phylogenomic analyses and molecular dating generated consistent evidence that supported the occurrence of a whole genome duplication event (termed ψ) before the species radiation within the BDTF clade (Bryidae, Dicranidae, Timmiidae and Funariidae) that is composed of the majority of lineages within the Bryopsida. By investigating the phylogenetic timing of the emergence of *P. patens* syntelog pairs from gene family phylogenies, we were able to recognize and circumscribe the WGD1 and WGD2 events in the ancestor of BDTF clade (the ψ event, ~87 mya) and Funarioideae (~65 mya), respectively. The age of the BDTF clade was also estimated to be around 80 mya from large-scale molecular evidence. In addition, we identified another two ancestral polyploidy events: the most ancient one in the ancestor of all mosses and the second that occurred before the separation of Polytrichopsida/Tetraphidopsida and Bryopsida (BPT clade). Molecular dating estimation analyses suggested that these two ancestral duplication events in mosses occurred around 329 and 182 mya, respectively.

## Acknowledgments

This work was financially supported by National Science Foundation (Collaborative Research: Dimensions: Grant Number 1638972 to MJO) and the NSFC-Xinjiang Key Project (Grant Number U1703233 to DZ). The School of Life Sciences of The Chinese University of Hong Kong provided the computational resources for this project.

## Author Contributions

BG conceived the study, performed the bioinformatic analyses and wrote the manuscript. MC, DZ and AJW contributed to the data analyses and result discussion. MJO and JZ contributed to the data interpretation, critically revised and improved the manuscript. All authors read and approved the manuscript.

## Supplementary Material

Supplementary tables and figures mentioned in the manuscript are available in the supplementary data.

## References

Arrigo N, Barker MS. 2012. Rarely successful polyploids and their legacy in plant genomes. Curr Opin Plant Biol 15(2): 140–146.

Bell NE, Quandt D, O’Brien TJ, Newton AE. 2007. Taxonomy and phylogeny in the earliest diverging pleurocarps: square holes and bifurcating pegs. Bryologist 110(3): 533–560.

Blanc G, Hokamp K, Wolfe KH. 2003. A recent polyploidy superimposed on older large-scale duplications in the Arabidopsis genome. Genome Res 13(2): 137–144.

Blanc G, Wolfe KH. 2004. Widespread Paleopolyploidy in Model Plant Species Inferred from Age Distributions of Duplicate Genes. Plant Cell 16(7): 1667–1678.

Bolger AM, Lohse M, Usadel B. 2014. Trimmomatic: a flexible trimmer for Illumina sequence data. Bioinformatics 30(15): 2114–2120.

Bowers JE, Chapman BA, Rong J, Paterson AH. 2003. Unravelling angiosperm genome evolution by phylogenetic analysis of chromosomal duplication events. Nature 422(6930): 433–438.

Buchfink B, Xie C, Huson DH. 2015. Fast and sensitive protein alignment using DIAMOND. Nat Methods 12(1): 59–60.

Capella-Gutierrez S, Silla-Martinez JM, Gabaldon T. 2009. trimAl: a tool for automated alignment trimming in large-scale phylogenetic analyses. Bioinformatics 25(15): 1972–1973.

Christiano De Souza IC, Ricardi Branco FS, Vargas YL. 2012. Permian bryophytes of Western Gondwanaland from the Paraná Basin in Brazil. Palaeontology 55(1): 229–241.

Cui L, Wall PK, Leebens-Mack JH, Lindsay BG, Soltis DE, Doyle JJ, Soltis PS, Carlson JE, Arumuganathan K, Barakat A, et al. 2006. Widespread genome duplications throughout the history of flowering plants. Genome Res 16(6): 738–749.

Devos N, Szovenyi P, Weston DJ, Rothfels CJ, Johnson MG, Shaw AJ. 2016. Analyses of transcriptome sequences reveal multiple ancient large-scale duplication events in the ancestor of Sphagnopsida (Bryophyta). New Phytol 211(1): 300–318.

Doyle JJ, Egan AN. 2010. Dating the origins of polyploidy events. New Phytol 186(1): 73–85.

Edgar RC. 2004. MUSCLE: multiple sequence alignment with high accuracy and high throughput. Nucleic Acids Res 32(5): 1792–1797.

Emms DM, Kelly S. 2015. OrthoFinder: solving fundamental biases in whole genome comparisons dramatically improves orthogroup inference accuracy. Genome Biol 16(1): 157.

Engel PP. 1968. The induction of biochemical and morphological mutants in the moss Physcomitrella patens. Am JBot 55(4): 438–446.

Fiz-Palacios O, Schneider H, Heinrichs J, Savolainen V. 2011. Diversification of land plants: insights from a family-level phylogenetic analysis. BMC Evol Biol 11: 341.

Friedman WE. 2009. The meaning of Darwin’s ‘abominable mystery’. Am J Bot 96(1): 5–21.

Fritsch R. 1991. Index to bryophyte chromosome counts: J. Cramer.

Gao B, Li X, Zhang D, Liang Y, Yang H, Chen M, Zhang Y, Zhang J, Wood AJ. 2017. Desiccation tolerance in bryophytes: The dehydration and rehydration transcriptomes in the desiccation-tolerant bryophyte Bryum argenteum. Sci Rep 7(1): 7571.

Gao B, Zhang D, Li X, Yang H, Wood AJ. 2014. De novo assembly and characterization of the transcriptome in the desiccation-tolerant moss Syntrichia caninervis. BMC Res Notes 7(1): 490.

Goffinet B. 2004. Systematics of the Bryophyta (mosses): from molecules to a revised classification. Molecular Systematics of Bryophytes. Monographs in Systematic Botany 98: 205–239.

Goldman N, Yang Z. 1994. A codon-based model of nucleotide substitution for protein-coding DNA sequences. Mol Biol Evol 11(5): 725–736.

Haas BJ, Papanicolaou A, Yassour M, Grabherr M, Blood PD, Bowden J, Couger MB, Eccles D, Li B, Lieber M, et al. 2013. De novo transcript sequence reconstruction from RNA-seq using the Trinity platform for reference generation and analysis. Nat Protoc 8(8): 1494–1512.

Ignatov MS. 1990. Upper Permian mosses from the Russian platform. Palaeontographica Abteilung B: 147–189.

Jaillon O, Aury JM, Noel B, Policriti A, Clepet C, Casagrande A, Choisne N, Aubourg S, Vitulo N, Jubin C, et al. 2007. The grapevine genome sequence suggests ancestral hexaploidization in major angiosperm phyla. Nature 449(7161): 463–467.

Jiao Y, Li J, Tang H, Paterson AH. 2014. Integrated syntenic and phylogenomic analyses reveal an ancient genome duplication in monocots. Plant Cell 26(7): 2792–2802.

Jiao Y, Wickett NJ, Ayyampalayam S, Chanderbali AS, Landherr L, Ralph PE, Tomsho LP, Hu Y, Liang H, Soltis PS, et al. 2011. Ancestral polyploidy in seed plants and angiosperms. Nature 473(7345): 97–100.

Jiao YN, Leebens-Mack J, Ayyampalayam S, Bowers JE, McKain MR, McNeal J, Rolf M, Ruzicka DR, Wafula E, Wickett NJ, et al. 2012. A genome triplication associated with early diversification of the core eudicots. Genome Biol 13(1): R3.

Johnson MG, Malley C, Goffinet B, Shaw AJ, Wickett NJ. 2016. A phylotranscriptomic analysis of gene family expansion and evolution in the largest order of pleurocarpous mosses (Hypnales, Bryophyta). Mol Phylogenet Evol 98: 29–40.

Katoh K, Standley DM. 2013. MAFFT multiple sequence alignment software version 7: improvements in performance and usability. Mol Biol Evol 30(4): 772–780.

Krzywinski M, Schein J, Birol I, Connors J, Gascoyne R, Horsman D, Jones SJ, Marra MA. 2009. Circos: an information aesthetic for comparative genomics. Genome Res 19(9): 1639–1645.

Laenen B, Shaw B, Schneider H, Goffinet B, Paradis E, Desamore A, Heinrichs J, Villarreal JC, Gradstein SR, McDaniel SF, et al. 2014. Extant diversity of bryophytes emerged from successive post-Mesozoic diversification bursts. Nat Commun 5: 5134.

Lanfear R, Frandsen PB, Wright AM, Senfeld T, Calcott B. 2017. PartitionFinder 2: New Methods for Selecting Partitioned Models of Evolution for Molecular and Morphological Phylogenetic Analyses. Mol Biol Evol 34(3): 772–773.

Lang D, Ullrich KK, Murat F, Fuchs J, Jenkins J, Haas FB, Piednoel M, Gundlach H, Van Bel M, Meyberg R, et al. 2018. The Physcomitrella patens chromosome-scale assembly reveals moss genome structure and evolution. Plant J 93(3): 515–533.

Larsén E, Rydin C. 2016. Disentangling the Phylogeny of Isoetes (Isoetales), Using Nuclear and Plastid Data. International journal of plant sciences 177(2): 157–174.

Li W, Godzik A. 2006. Cd-hit: a fast program for clustering and comparing large sets of protein or nucleotide sequences. Bioinformatics 22(13): 1658–1659.

Li Z, Baniaga AE, Sessa EB, Scascitelli M, Graham SW, Rieseberg LH, Barker MS. 2015. Early genome duplications in conifers and other seed plants. Sci Adv 1(10): e1501084.

Lorrain S, Lin B, Auriac MC, Kroj T, Saindrenan P, Nicole M, Balague C, Roby D. 2004. Vascular associated death1, a novel GRAM domain-containing protein, is a regulator of cell death and defense responses in vascular tissues. Plant Cell 16(8): 2217–2232.

Magallon S, Hilu KW, Quandt D. 2013. Land plant evolutionary timeline: gene effects are secondary to fossil constraints in relaxed clock estimation of age and substitution rates. Am J Bot 100(3): 556–573.

Mayrose I, Zhan SH, Rothfels CJ, Magnuson-Ford K, Barker MS, Rieseberg LH, Otto SP. 2011. Recently formed polyploid plants diversify at lower rates. Science 333(6047): 1257.

McKain MR, Tang H, McNeal JR, Ayyampalayam S, Davis JI, dePamphilis CW, Givnish TJ, Pires JC, Stevenson DW, Leebens-Mack JH. 2016. A Phylogenomic Assessment of Ancient Polyploidy and Genome Evolution across the Poales. Genome Biol Evol 8(4): 1150–1164.

McLachlan GJ, Peel D, Basford KE, Adams P. 1999. The EMMIX software for the fitting of mixtures of normal and t-components. Journal of Statistical Software 4(2): 1–14.

Medina R, Johnson M, Liu Y, Wilding N, Hedderson TA, Wickett N, Goffinet B. 2018. Evolutionary dynamism in bryophytes: Phylogenomic inferences confirm rapid radiation in the moss family Funariaceae. Mol Phylogenet Evol 120: 240–247.

Morris JL, Puttick MN, Clark JW, Edwards D, Kenrick P, Pressel S, Wellman CH, Yang Z, Schneider H, Donoghue PCJ. 2018. The timescale of early land plant evolution. Proc Natl Acad Sci U S A 115(10): E2274–E2283.

Morrison DA. 2008. How to Summarize Estimates of Ancestral Divergence Times. Evolutionary Bioinformatics 4: 75–95.

Neuburg M. 1960. Leafy mosses from the Permian deposits of Angaraland. Trudy Geol. Inst. Akad. Nauk SSSR 19: 1–104.

Newton AE, Wikström N, Bell N, Lowe Forrest L, Ignatov MS. 2006. Dating the diversification of pleurocarpous mosses.

Nickrent DL, Parkinson CL, Palmer JD, Duff RJ. 2000. Multigene phylogeny of land plants with special reference to bryophytes and the earliest land plants. Mol Biol Evol 17(12): 1885–1895.

Ohno S. 1970. Evolution by Gene Duplication. (Berlin: Springer).

Price MN, Dehal PS, Arkin AP. 2010. FastTree 2-Approximately Maximum-Likelihood Trees for Large Alignments. PLoS ONE 5(3).

Qiu YL, Li L, Wang B, Chen Z, Knoop V, Groth-Malonek M, Dombrovska O, Lee J, Kent L, Rest J, et al. 2006. The deepest divergences in land plants inferred from phylogenomic evidence. Proc Natl Acad Sci U S A 103(42): 15511–15516.

Ran JH, Shen TT, Wang MM, Wang XQ. 2018. Phylogenomics resolves the deep phylogeny of seed plants and indicates partial convergent or homoplastic evolution between Gnetales and angiosperms. Proc Biol Sci 285(1881).

Rensing SA, Ick J, Fawcett JA, Lang D, Zimmer A, Van de Peer Y, Reski R. 2007. An ancient genome duplication contributed to the abundance of metabolic genes in the moss Physcomitrella patens. BMC Evol Biol 7: 130.

Rensing SA, Lang D, Zimmer AD, Terry A, Salamov A, Shapiro H, Nishiyama T, Perroud PF, Lindquist EA, Kamisugi Y, et al. 2008. The Physcomitrella genome reveals evolutionary insights into the conquest of land by plants. Science 319(5859): 64–69.

Robertson FM, Gundappa MK, Grammes F, Hvidsten TR, Redmond AK, Lien S, Martin SAM, Holland PWH, Sandve SR, Macqueen DJ. 2017. Lineage-specific rediploidization is a mechanism to explain time-lags between genome duplication and evolutionary diversification. Genome Biol 18(1): 111.

Ruprecht C, Lohaus R, Vanneste K, Mutwil M, Nikoloski Z, Van de Peer Y, Persson S. 2017. Revisiting ancestral polyploidy in plants. Sci Adv 3(7): e1603195.

Salse J, Abrouk M, Bolot S, Guilhot N, Courcelle E, Faraut T, Waugh R, Close TJ, Messing J, Feuillet C. 2009. Reconstruction of monocotelydoneous proto-chromosomes reveals faster evolution in plants than in animals. Proc Natl Acad Sci U S A 106(35): 14908–14913.

Sanderson MJ. 2003. r8s: inferring absolute rates of molecular evolution and divergence times in the absence of a molecular clock. Bioinformatics 19(2): 301–302.

Schranz ME, Mohammadin S, Edger PP. 2012. Ancient whole genome duplications, novelty and diversification: the WGD Radiation Lag-Time Model. Curr Opin Plant Biol 15(2): 147–153.

Schween G, Schulte J, Reski R, Hohe A. 2005. Effect of ploidy level on growth, differentiation, and morphology in Physcomitrella patens. Bryologist 108(1): 27–35.

Shaw AJ, Cox CJ, Buck WR, Devos N, Buchanan AM, Cave L, Seppelt R, Shaw B, Larrain J, Andrus R, et al. 2010a. Newly resolved relationships in an early land plant lineage: Bryophyta class Sphagnopsida (peat mosses). Am J Bot 97(9): 1511–1531.

Shaw AJ, Devos N, Cox CJ, Boles SB, Shaw B, Buchanan AM, Cave L, Seppelt R. 2010b. Peatmoss (Sphagnum) diversification associated with Miocene Northern Hemisphere climatic cooling? Mol Phylogenet Evol 55(3): 1139–1145.

Shaw AJ, Szovenyi P, Shaw B. 2011. Bryophyte diversity and evolution: windows into the early evolution of land plants. Am J Bot 98(3): 352–369.

Shelton GWK, Stockey RA, Rothwell GW, Tomescu AMF. 2015. Exploring the fossil history of pleurocarpous mosses: Tricostaceae fam. nov from the Cretaceous of Vancouver Island, Canada. Am J Bot 102(11): 1883–1900.

Smith SA, Donoghue MJ. 2008. Rates of molecular evolution are linked to life history in flowering plants. Science 322(5898): 86–89.

Smith SA, Dunn CW. 2008. Phyutility: a phyloinformatics tool for trees, alignments and molecular data. Bioinformatics 24(5): 715–716.

Stamatakis A. 2014. RAxML version 8: a tool for phylogenetic analysis and post-analysis of large phylogenies. Bioinformatics 30(9): 1312–1313.

Suyama M, Torrents D, Bork P. 2006. PAL2NAL: robust conversion of protein sequence alignments into the corresponding codon alignments. Nucleic Acids Res 34(Web Server issue): W609–612.

Szovenyi P, Perroud PF, Symeonidi A, Stevenson S, Quatrano RS, Rensing SA, Cuming AC, McDaniel SF. 2015. De novo assembly and comparative analysis of the Ceratodon purpureus transcriptome. Mol Ecol Resour 15(1): 203215.

Tang HB, Wang XY, Bowers JE, Ming R, Alam M, Paterson AH. 2008. Unraveling ancient hexaploidy through multiply-aligned angiosperm gene maps. Genome Research 18(12): 1944–1954.

Unruh SA, McKain MR, Lee YI, Yukawa T, McCormick MK, Shefferson RP, Smithson A, Leebens-Mack JH, Pires JC. 2018. Phylotranscriptomic analysis and genome evolution of the Cypripedioideae (Orchidaceae). Am J Bot 105(4): 631–640.

Van de Peer Y, Mizrachi E, Marchal K. 2017. The evolutionary significance of polyploidy. Nat Rev Genet 18(7): 411–424.

Vanneste K, Baele G, Maere S, Van de Peer Y. 2014. Analysis of 41 plant genomes supports a wave of successful genome duplications in association with the Cretaceous-Paleogene boundary. Genome Res 24(8): 1334–1347.

Vekemans D, Proost S, Vanneste K, Coenen H, Viaene T, Ruelens P, Maere S, Van de Peer Y, Geuten K. 2012. Gamma paleohexaploidy in the stem lineage of core eudicots: significance for MADS-box gene and species diversification. Mol Biol Evol 29(12): 3793–3806.

Wang D, Zhang Y, Zhang Z, Zhu J, Yu J. 2010. KaKs_Calculator 2.0: a toolkit incorporating gamma-series methods and sliding window strategies. Genomics Proteomics Bioinformatics 8(1): 77–80.

Wang Y, Tang H, Debarry JD, Tan X, Li J, Wang X, Lee TH, Jin H, Marler B, Guo H, et al. 2012. MCScanX: a toolkit for detection and evolutionary analysis of gene synteny and collinearity. Nucleic Acids Res 40(7): e49.

Wickett NJ, Mirarab S, Nguyen N, Warnow T, Carpenter E, Matasci N, Ayyampalayam S, Barker MS, Burleigh JG, Gitzendanner MA, et al. 2014. Phylotranscriptomic analysis of the origin and early diversification of land plants. Proc Natl Acad Sci U S A 111(45): E4859–4868.

Yang Y, Moore MJ, Brockington SF, Mikenas J, Olivieri J, Walker JF, Smith SA. 2018. Improved transcriptome sampling pinpoints 26 ancient and more recent polyploidy events in Caryophyllales, including two allopolyploidy events. New Phytol 217(2): 855–870.

Yang Y, Smith SA. 2014. Orthology Inference in Nonmodel Organisms Using Transcriptomes and Low-Coverage Genomes: Improving Accuracy and Matrix Occupancy for Phylogenomics. Molecular Biology and Evolution 31(11): 3081–3092.

Zhang C, Rabiee M, Sayyari E, Mirarab S. 2018. ASTRAL-III: polynomial time species tree reconstruction from partially resolved gene trees. BMC Bioinformatics 19(Suppl 6): 153.

