## Supplementary materials for "Integrated phylogenomic analyses reveal recurrent ancestral large-scale duplication events in mosses"

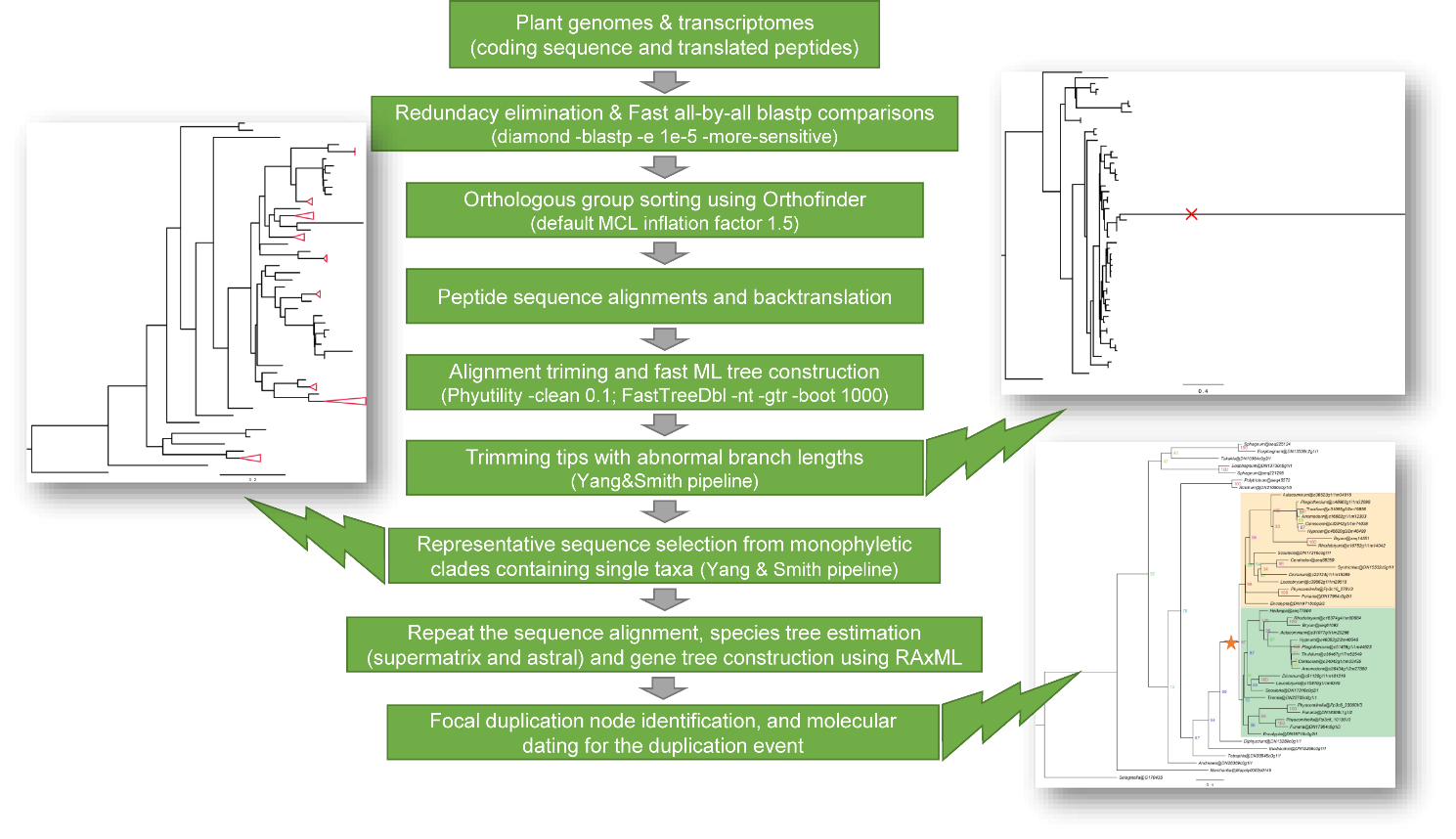

**Figure S1 – Phylogenomic workflow utilized in this study for species tree construction and identification of ancestral gene duplications in mosses.**

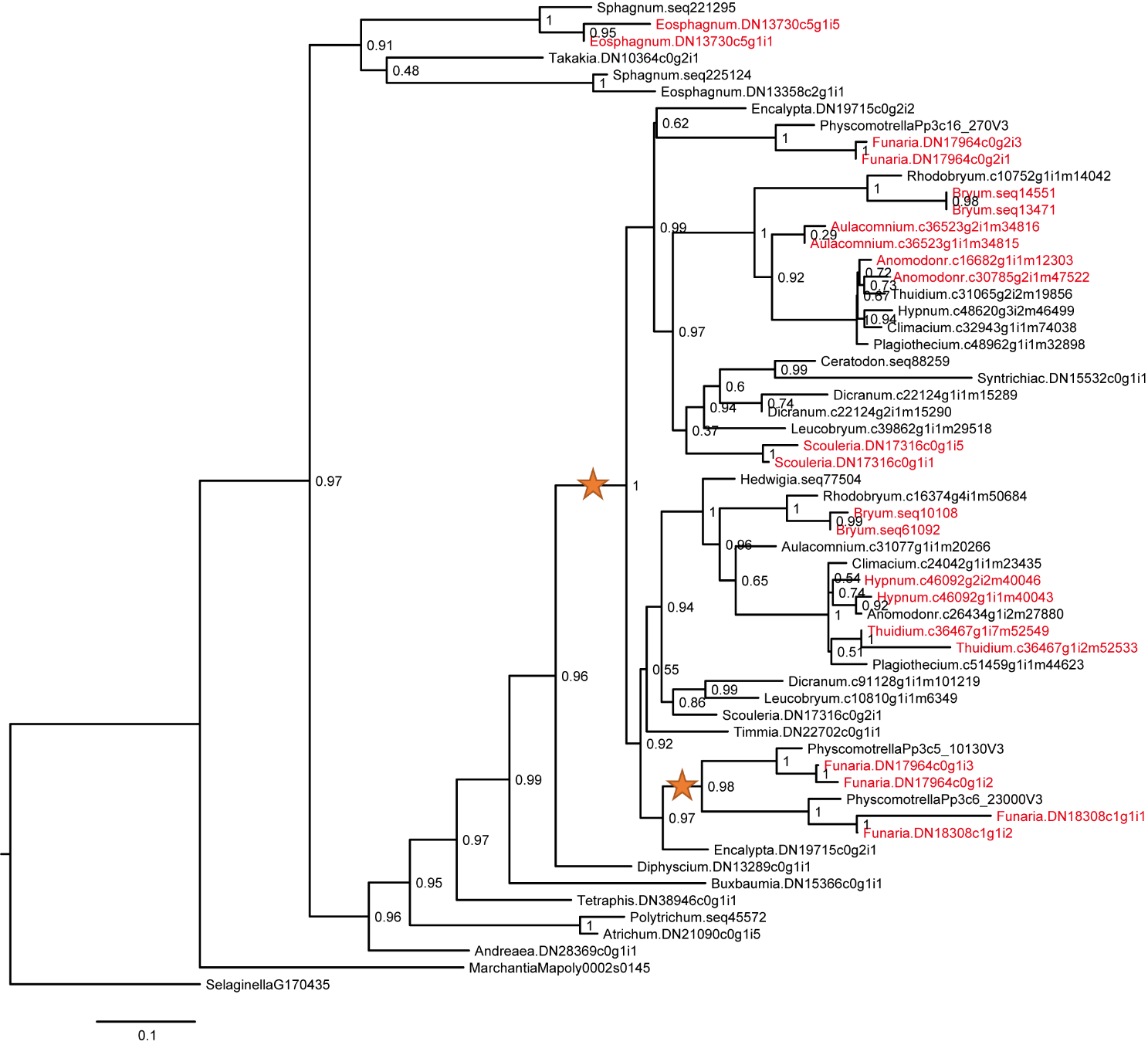

**Figure S2 – Exemplary gene tree generated in FastTree before masking the single-taxa monophyletic and paraphyletic clades.** Clades that were constituted with genes from a single species were highlighted in red and the two ancestral duplications in Funarioideae and BDTF clade were labeled with stars.

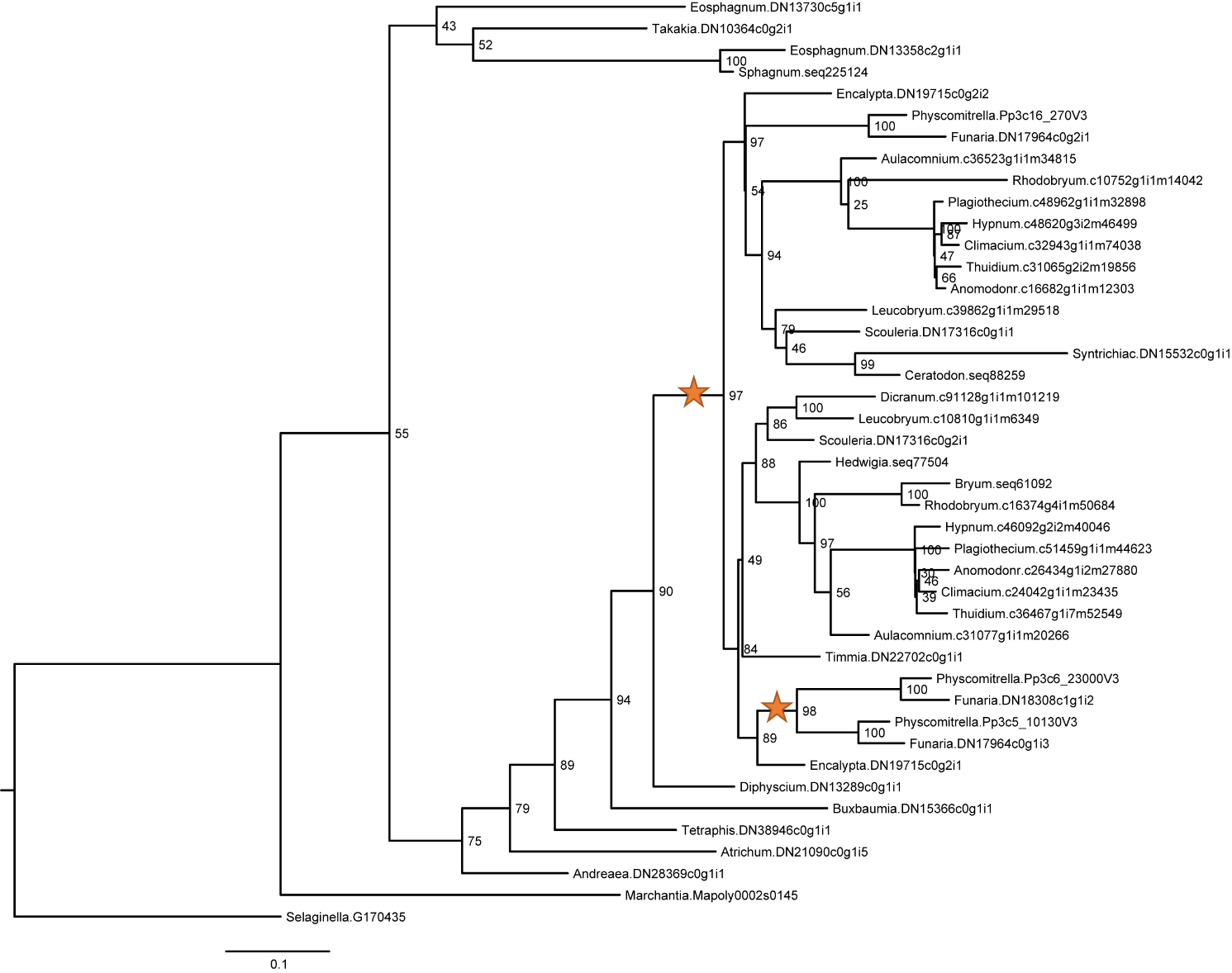

**Figure S3 – Exemplary gene tree generated in RAxML after masking the single-taxa monophyletic and paraphyletic clades.** Clades constituted with genes from a single species in Figure S2 were masked and one representative sequence retained using the Yang & Smith (2014) pipeline. The detection of the two ancestral duplications in Funarioideae and BDTF (labeled with stars) were not affected by this masking procedure.

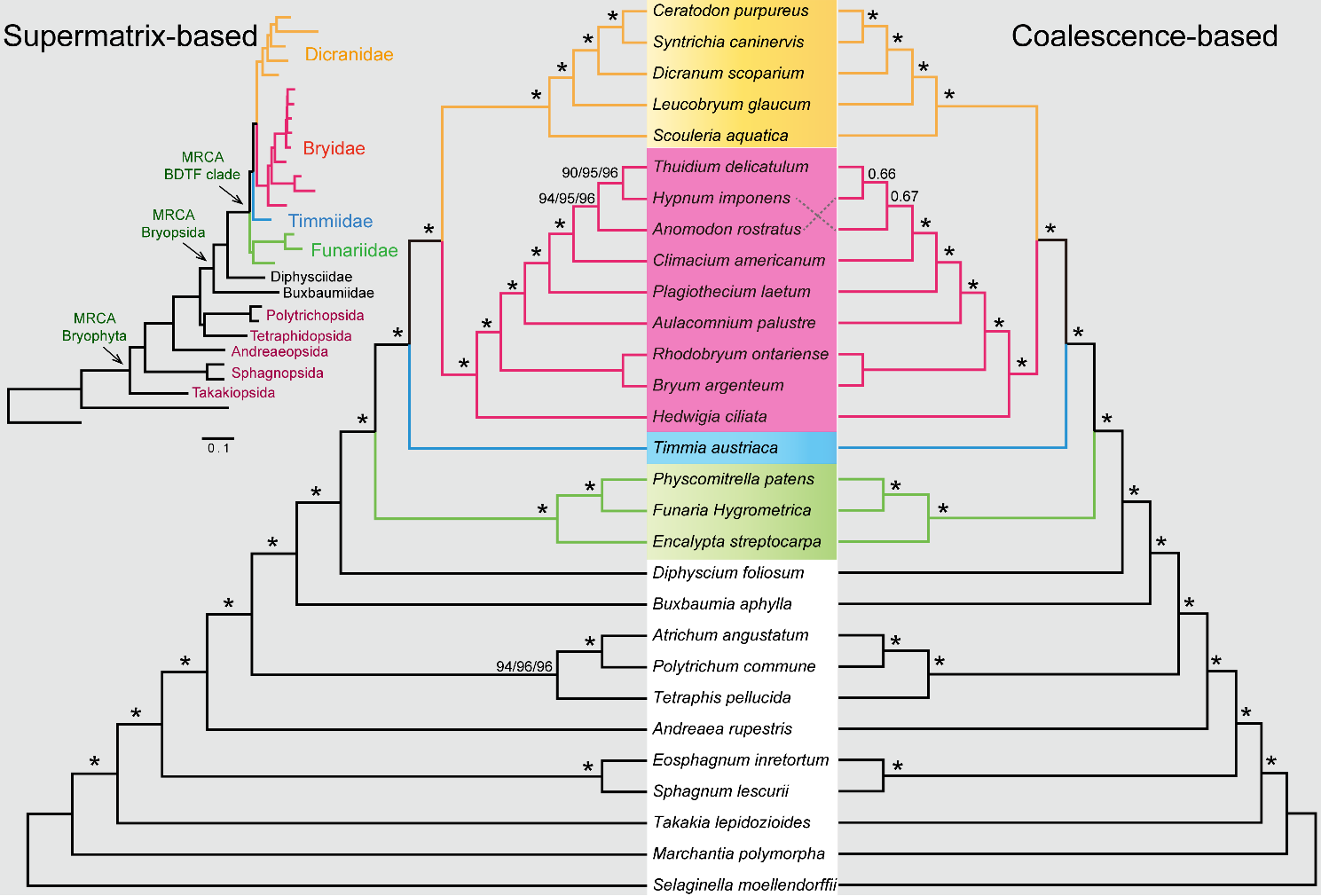

**Figure S4 – Comparison of species trees constructed using supermatrix-based and coalescence-based approaches.** Left panel depicts the maximum-likelihood nucleotide supermatrix tree topology from RAxML. The right panel depicts the coalescence-based species tree inferred from ASTRAL-III, generated from the 649 maximum-likelihood gene trees of each single-copy orthogroup. Nodal bootstrap support values indicated by asterisks are 100%. The supermatrix-based and coalescence-based approaches generated consistent species tree topology at the subclass level in mosses, the only variation was discovered in the clade of Hypnales.

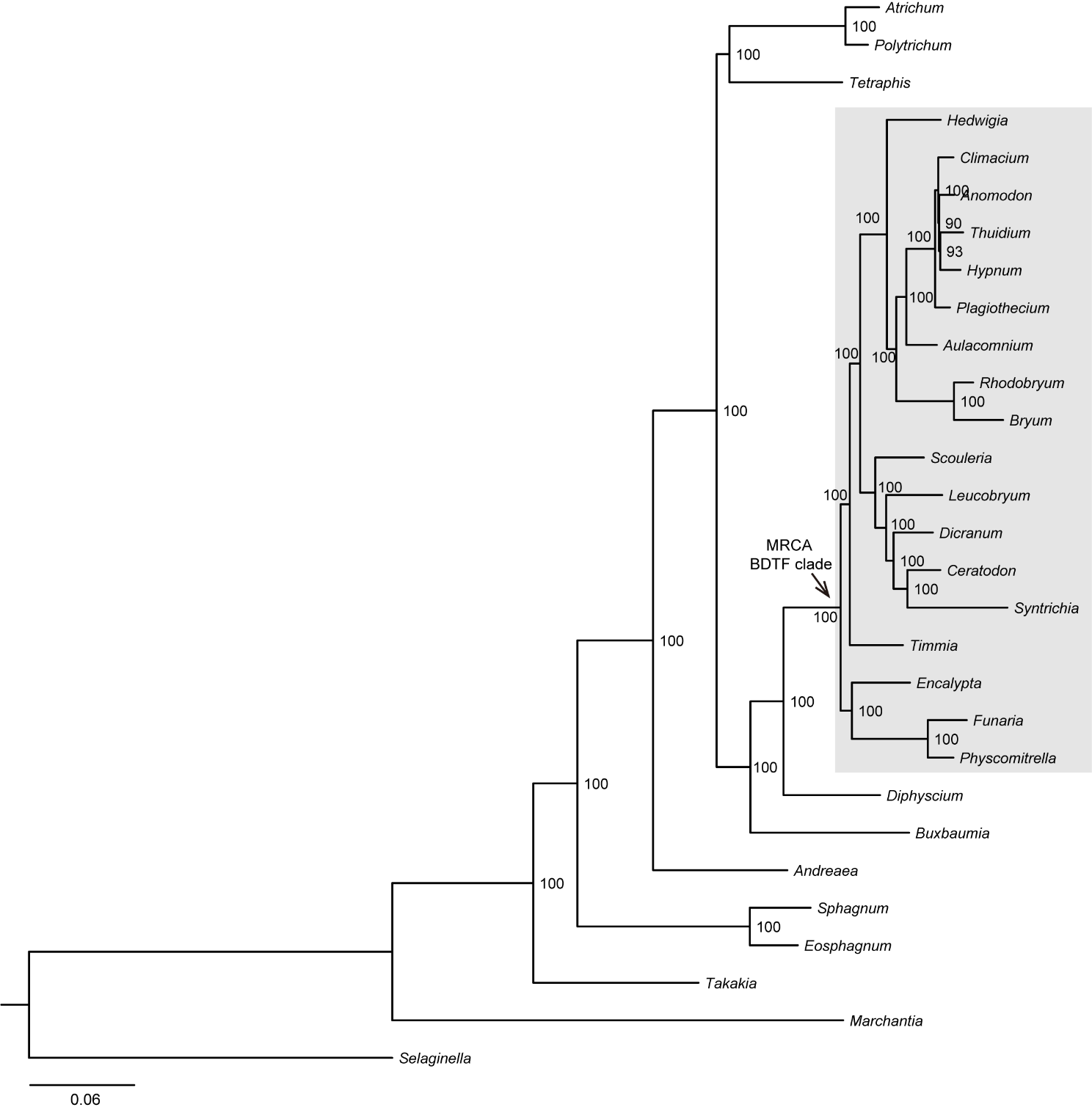

**Figure S5 – The maximum-likelihood species tree generated using the 1st and 2nd codon positions of the concatenated alignment supermatrix.** Branch lengths are scaled to substitutions per site. The MRCA node for BDTF clade was indicated, and the relatively short branch length might be congruent with the rapid speciation following the ψ WGD event.

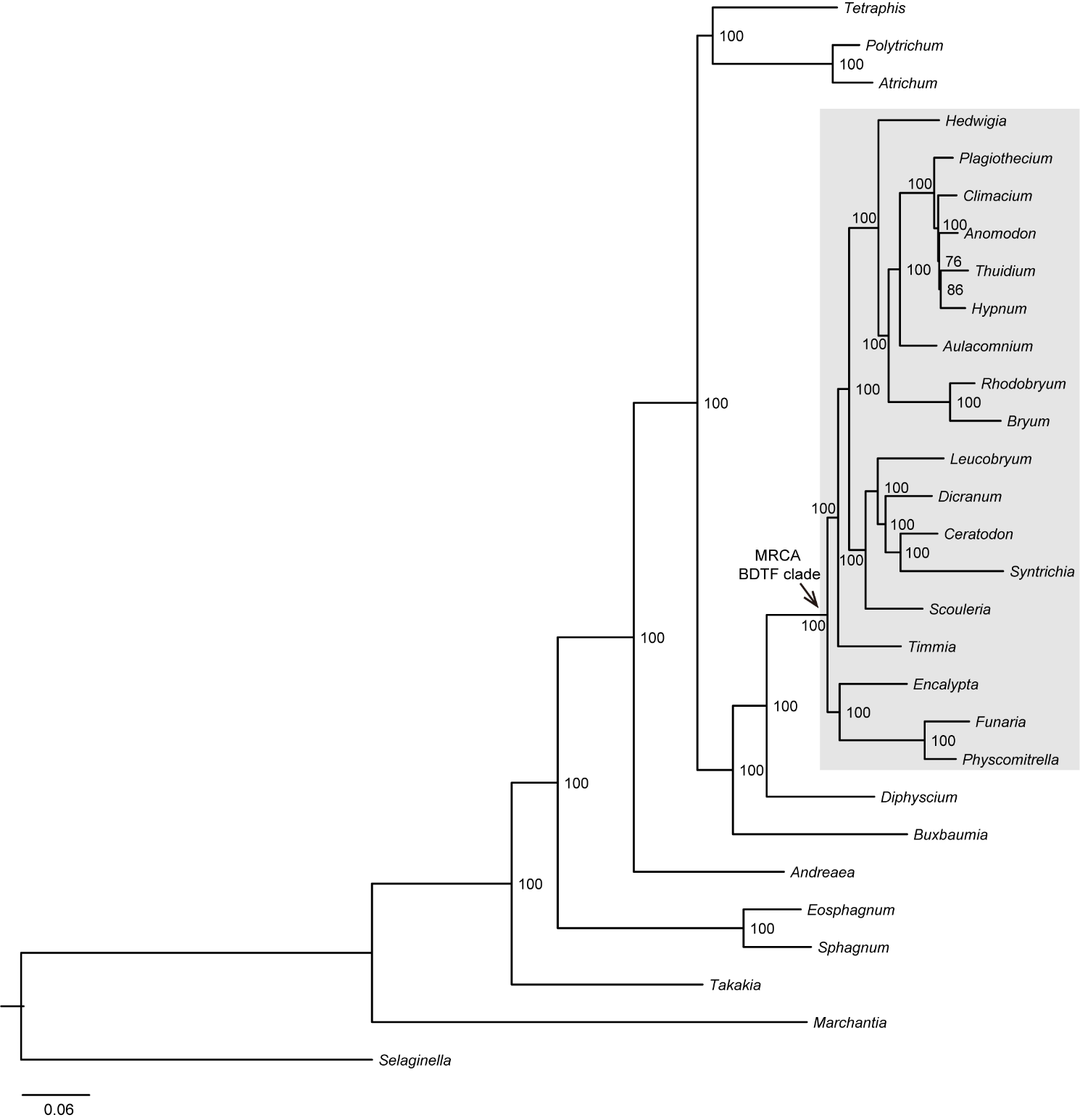

**Figure S6 – The maximum-likelihood species tree generated from the concatenated protein alignment using RAxML under the PROTGAMMAAUTO model.** Branch lengths are scaled to substitutions per site. The MRCA node for BDTF clade was indicated and highlighted.

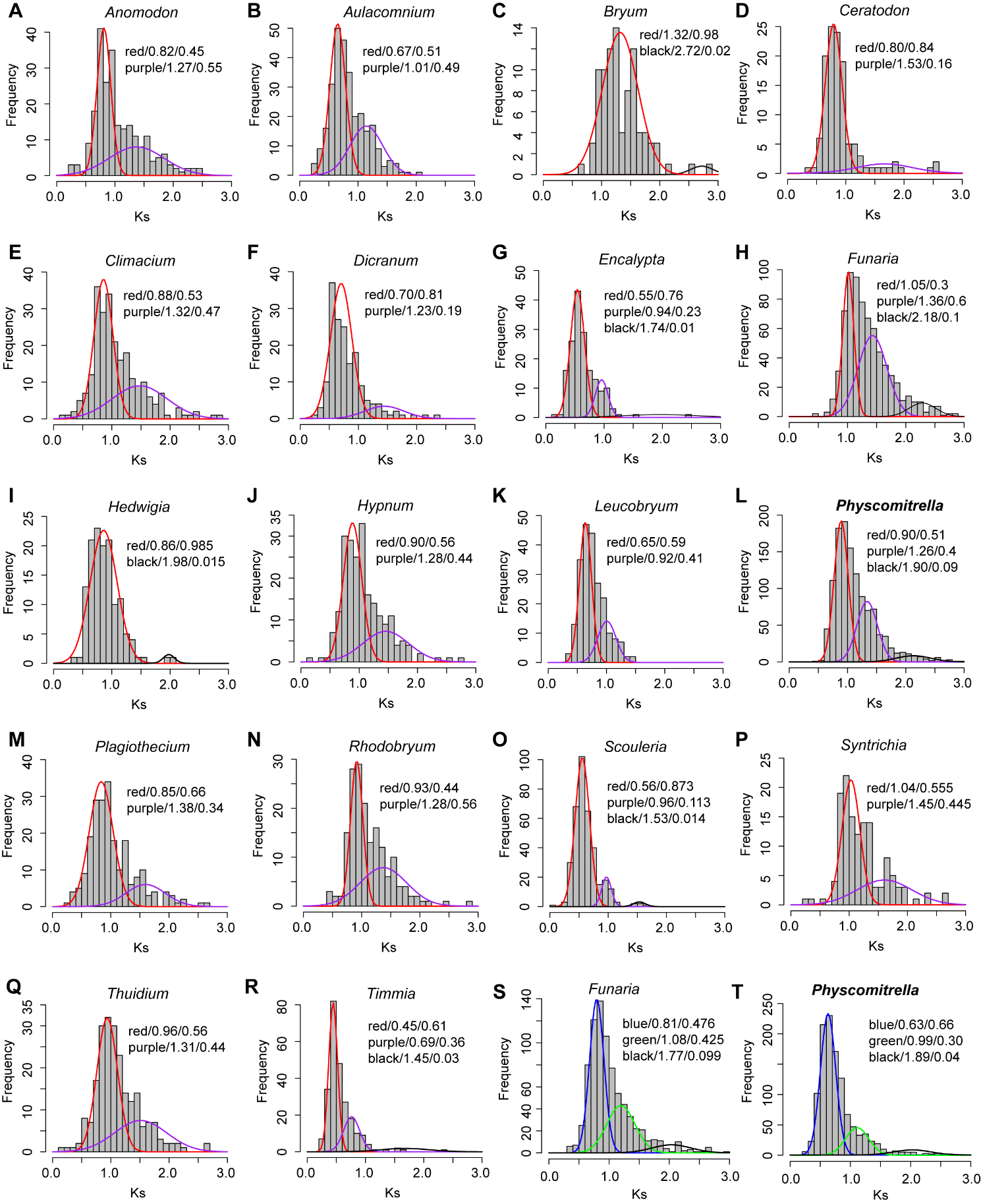

**Figure S7 - *Ks* distribution of all inferred duplicated paralogs in different species tracing the ψ and Funarioideae duplications. (A-S)** Paralogous pairs with coalescence node at N20 (BSV ≥ 50) were identified in each BDTF clade species and corresponding *Ks* distributions were analyzed. **(S-T)** *Ks* distribution analyses of paralogous pairs with coalescence node at N4 (BSV ≥ 50) in *Funaria hygrometrica* and *Physcomitrella patens*.

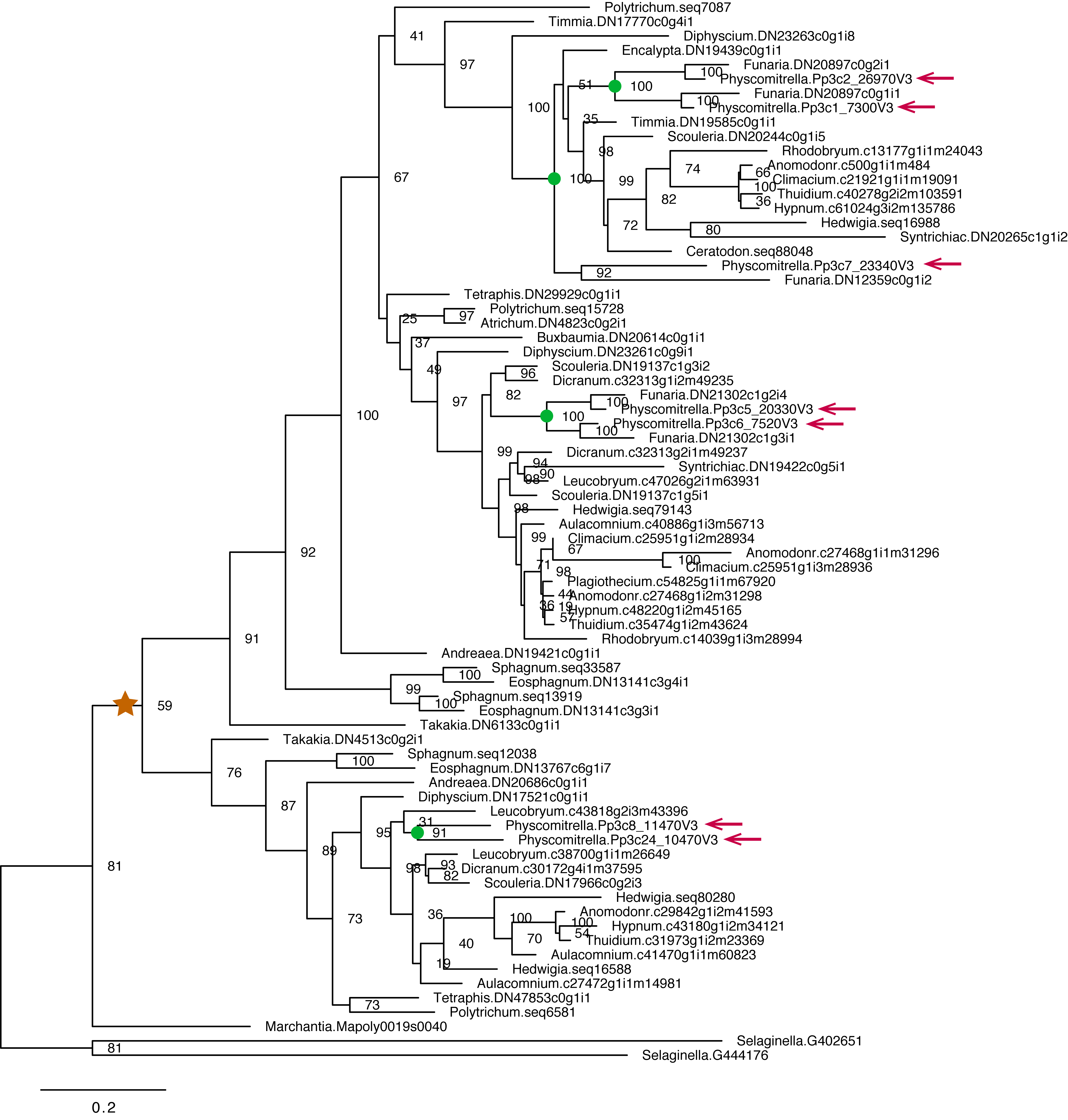

**Figure S8 - Exemplar maximum-likelihood gene family (OG-243) phylogeny contains *Physcomitrella patens* paralogs that survived ancestral duplication.** Genes in orthogroup_243 encode the bZIP transcription factors. *P. patens* genes were highlighted with arrows and the moss-wide duplication node was labeled as brown star. More recent duplications (e.g. the ψ and Funarioideae duplications) supported by tree topology were labeled as green circles.

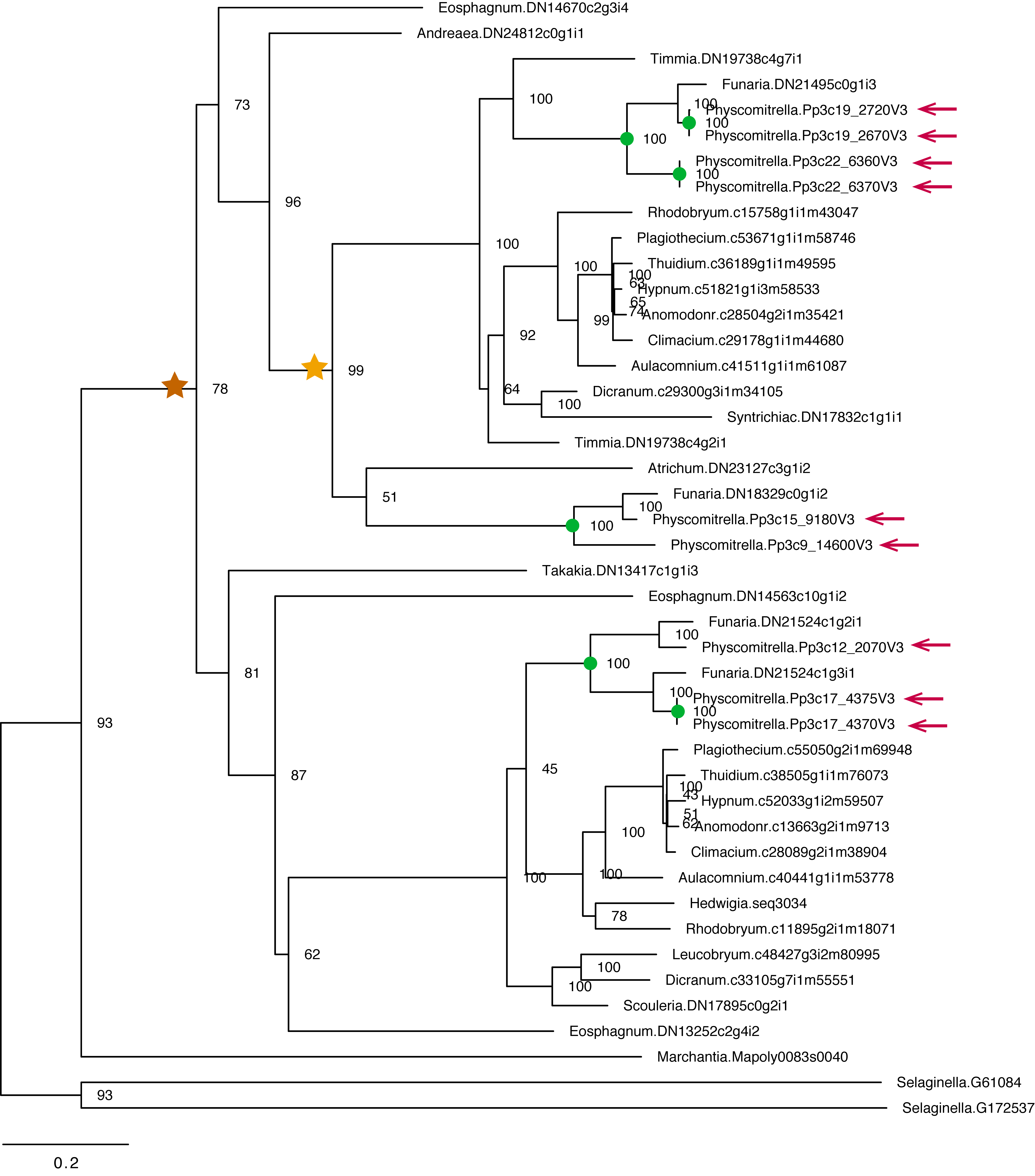

**Figure S9 - Exemplar maximum-likelihood gene family (OG-362) phylogeny contains *Physcomitrella patens* paralogs that survived ancestral duplications.** Genes in orthogroup_362 encode the Nin-like transcription factors. *P. patens* genes were highlighted with arrows and the moss-wide duplication node was labeled as brown star and the BPT duplication was labeled as a yellow star. More recent duplications supported by tree topology were labeled as green circles.

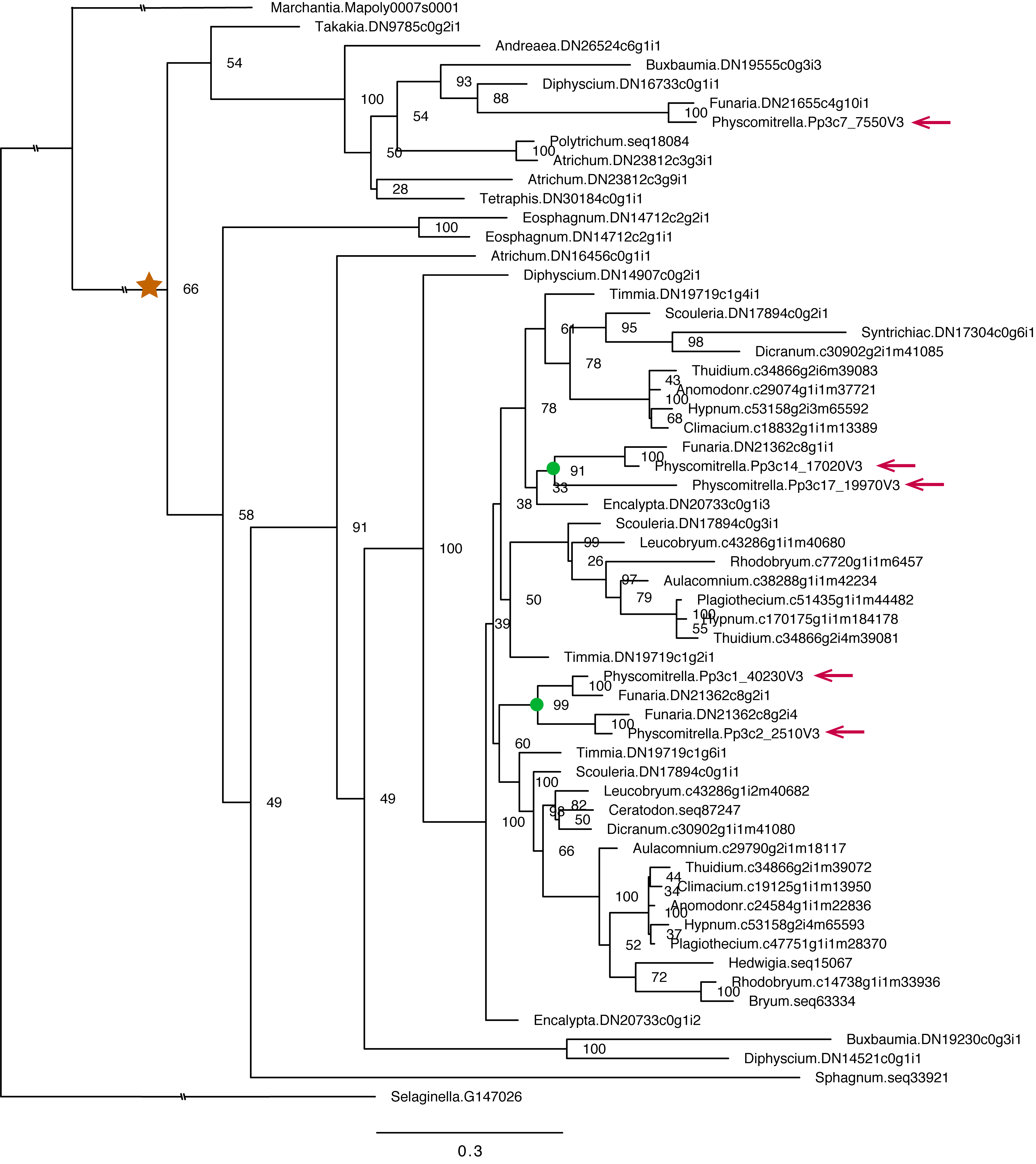

**Figure S10 - Exemplar maximum-likelihood gene family (OG-504) phylogeny contains *Physcomitrella patens* paralogs that survived ancestral duplications.** Genes in orthogroup_504 encode the WRKY transcription factors. *P. patens* genes were highlighted with arrows and the moss-wide duplication node was labeled as brown star. More recent duplications supported by tree topology were labeled as green circles.

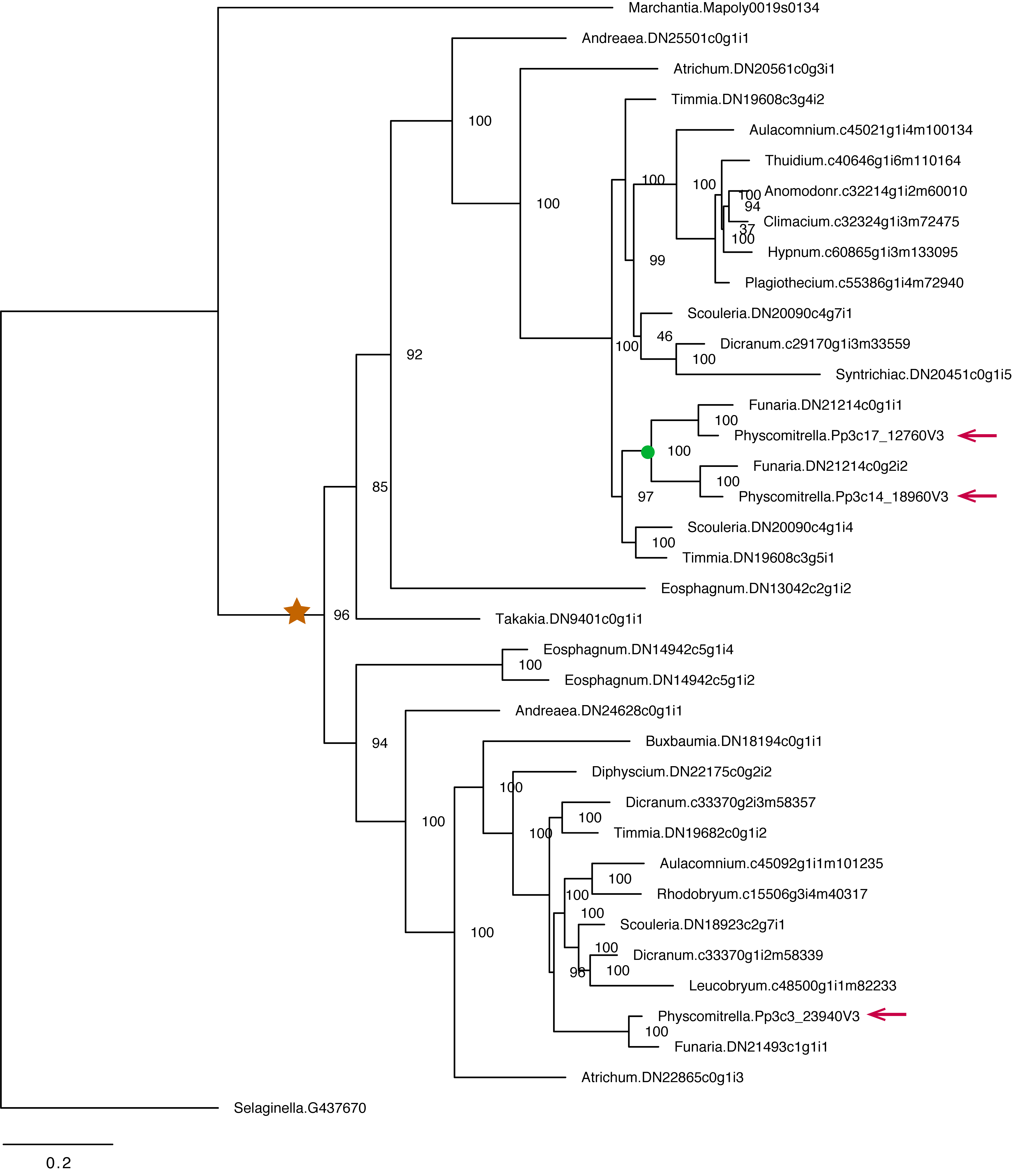

**Figure S11 - Exemplar maximum-likelihood gene family (OG-1137) phylogeny** **contains *Physcomitrella patens* paralogs that survived ancestral duplications.** Genes in orthogroup_1137 encode the SBP transcription factors. *P. patens* genes were highlighted with arrows and the moss-wide duplication node was labeled as brown star. More recent Funarioideae duplication supported by tree topology was highlighted with green circle.

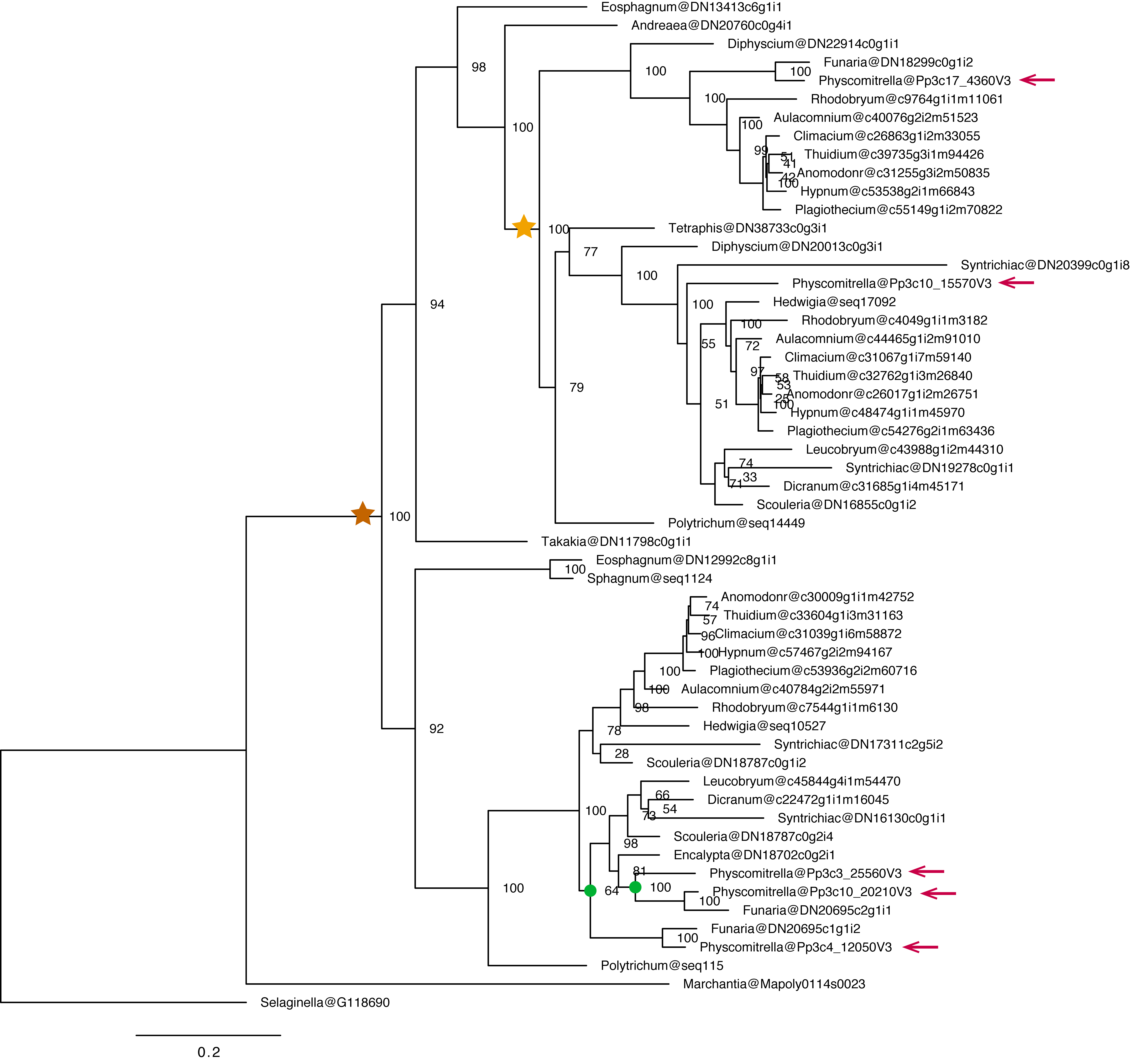

**Figure S12 - An exemplar maximum likelihood gene tree from** **OrthoGroup_462 contains *Physcomitrella patens* paralogs that survived ancestral duplications.** Genes in orthogroup_462 encodes homologs of VAD1 (Vascular Associated Death1), a regulator of cell death and defense responses in vascular tissues. *P. patens* genes were highlighted with arrows and the moss-wide duplication node was labeled as brown star and the BPT duplication was labeled as a yellow star. More recent *P. patens* gene duplications supported by tree topology were labeled as green circles.

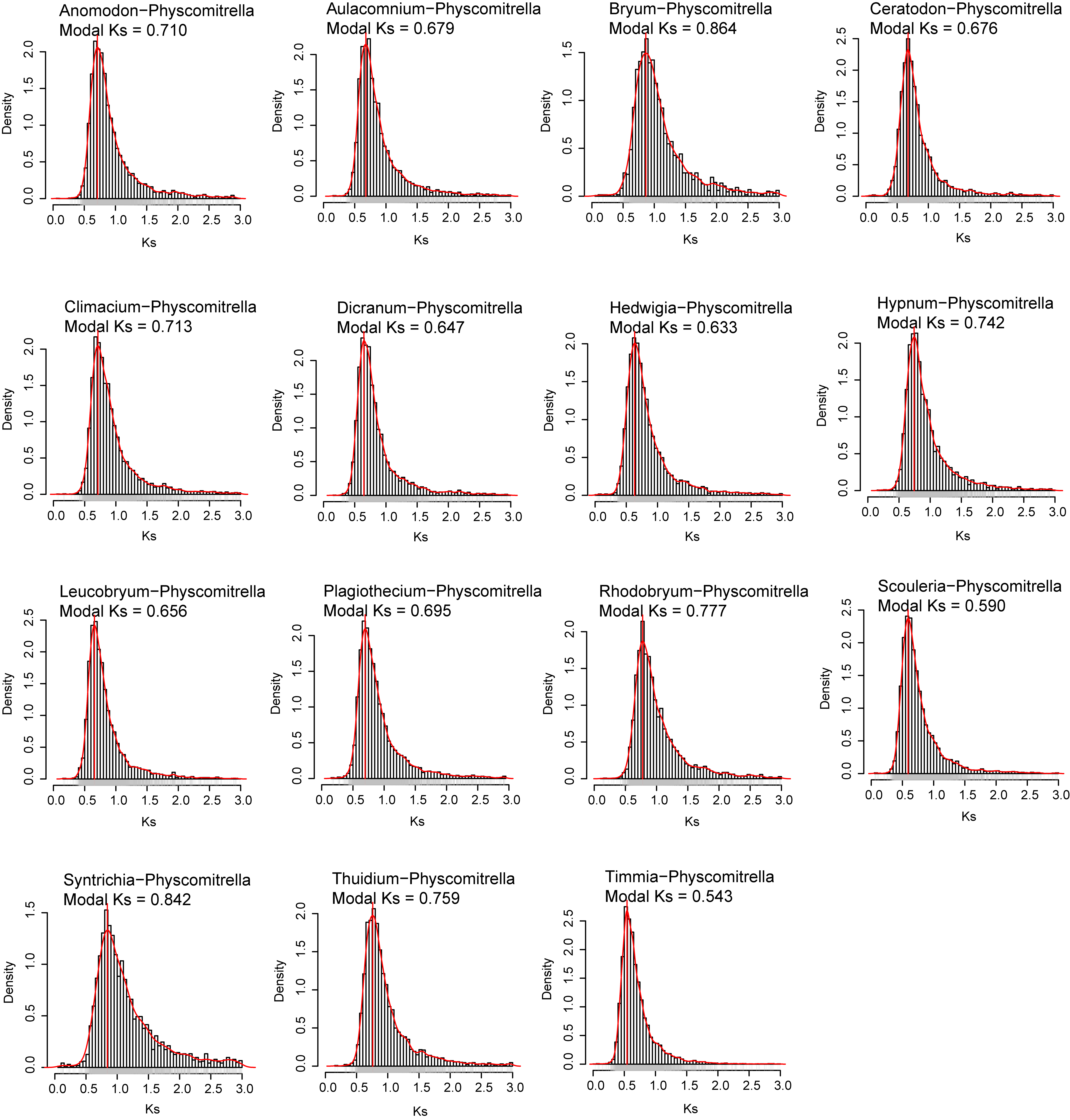

**Figure S13 -** Distribution of synonymous substitutions (Ks) in orthologous pairs between species of Bryidae, Dicranidae, Timmiidae and Physcomitrella patens. The modal Ks values were calculated from Kernal Density Estimation function in R statistical environment and labeled in the orthologous Ks distribution, providing supporting evidence for the relatively young age of the BDTF clade.

**Table S1 - Summary of plant genomes and transcriptomes included in the phylogenomic analyses**

| **Species** | **Lineage** | **Data source** | **Ref** | **Accession** | **Non-redundant peptides** |
| --- | --- | --- | --- | --- | --- |
| *Ceratodon purpureus* | Dicranidae | 1KP-pilot | [1] | ERR364350 | 17,420 |
| *Syntrichia caninervis** | Dicranidae | NCBI-SRA | [2] | SRR7345801 | 28,957 |
| *Dicranum scoparium* | Dicranidae | DRYAD | [3] | SRR2518102 | 13,368 |
| *Leucobryum glaucum* | Dicranidae | DRYAD | [3] | SRR2518106 | 12,399 |
| *Scouleria aquatica** | Dicranidae | 1KP | [4] | ERR2040953 | 31,394 |
| *Thuidium delicatulum* | Bryidae-Hypnales | DRYAD | [3] | SRR2499574 | 19,883 |
| *Hypnum imponens* | Bryidae-Hypnales | DRYAD | [3] | SRR2513252 | 21,509 |
| *Anomodon rostratus* | Bryidae-Hypnales | DRYAD | [3] | SRR2518087 | 19,435 |
| *Climacium americanum* | Bryidae-Hypnales | DRYAD | [3] | SRR2497426 | 19,642 |
| *Plagiothecium laetum* | Bryidae-Hypnales | DRYAD | [3] | SRR2518105 | 20,315 |
| *Aulacomnium palustre* | Bryidae | DRYAD | [3] | SRR2518059 | 17,878 |
| *Rhodobryum ontariense* | Bryidae | DRYAD | [3] | SRR2518100 | 11,140 |
| *Bryum argenteum* | Bryidae | 1KP-pilot | [1] | ERR364348 | 14,458 |
| *Hedwigia ciliata* | Bryidae | 1KP-pilot | [1] | ERR364352 | 16,853 |
| *Timmia austriaca** | Timmiidae | 1KP | [4] | ERR2040975 | 23,851 |
| *Physcomitrella patens* | Funariidae-Funariaceae | Genome v3.3 | [4] | Phytozome 12.1.6 | 32,926 |
| *Funaria Hygrometrica** | Funariidae-Funariaceae | NCBI-SRA | [5] | SRR6356272 | 27,978 |
| *Encalypta streptocarpa** | Funariidae-Encalyptaceae | 1KP | [4] | ERR2040959 | 21,787 |
| *Diphyscium foliosum** | Diphysciidae | 1KP | [4] | ERR2040958 | 23,398 |
| *Buxbaumia aphylla** | Buxbaumiidae | 1KP | [4] | ERR2040954 | 21,646 |
| *Atrichum angustatum** | Polytrichopsida | 1KP | [4] | ERR2040969 | 26,405 |
| *Polytrichum commune* | Polytrichopsida | 1KP-pilot | [1] | ERR364413 | 11,193 |
| *Tetraphis pellucida** | Tetraphidopsida | 1KP | [4] | ERR2040974 | 28,215 |
| *Andreaea rupestris** | Andreaeopsida | 1KP | [4] | ERR2040947 | 24,498 |
| *Eosphagnum inretortum** | Sphagnopsida | NCBI-SRA | [6] | ERR2324899-  ERR2324902 | 45,329 |
| *Sphagnum lescurii* | Sphagnopsida | 1KP-pilot | [1] | ERR364414 | 26,160 |
| *Takakia lepidozioides** | Takakiopsida | 1KP | [4] | ERR2040973 | 23,411 |
| *Marchantia polymorpha* | Liverwort | Genome v3.1 | [7] | Phytozome 12.1.6 | 19,287 |
| *Selaginella moellendorffii* | Tracheophyte | Genome v1.0 | [8] | Phytozome 12.1.6 | 22,285 |

*New Trinity transcriptome assemblies generated from reads publicly deposited in NCBI-SRA database.

**[1] Wickett NJ, Mirarab S, Nguyen N, Warnow T, Carpenter E, Matasci N, Ayyampalayam S, Barker MS, Burleigh JG, Gitzendanner MA, et al. 2014.** Phylotranscriptomic analysis of the origin and early diversification of land plants. *Proceedings of the National Academy of Sciences* **111**(45): E4859-E4868.

**[2] Gao B, Zhang D, Li X, Yang H, Wood AJ. 2014.** De novo assembly and characterization of the transcriptome in the desiccation-tolerant moss Syntrichia caninervis. *BMC Res Notes* **7**(1): 490.

**[3] Johnson MG, Malley C, Goffinet B, Shaw AJ, Wickett NJ. 2016.** A phylotranscriptomic analysis of gene family expansion and evolution in the largest order of pleurocarpous mosses (Hypnales, Bryophyta). *Molecular Phylogenetics and Evolution* **98**: 29-40.

**[4] Lang D, Ullrich KK, Murat F, Fuchs J, Jenkins J, Haas FB, Piednoel M, Gundlach H, Van Bel M, Meyberg R, et al. 2018.** The Physcomitrella patens chromosome-scale assembly reveals moss genome structure and evolution. *The Plant Journal* **93**(3): 515-533.

**[5] Medina R, Johnson M, Liu Y, Wilding N, Hedderson TA, Wickett N, Goffinet B. 2018.** Evolutionary dynamism in bryophytes: Phylogenomic inferences confirm rapid radiation in the moss family Funariaceae. *Mol Phylogenet Evol* **120**: 240-247.

**[6] Devos N, Szovenyi P, Weston DJ, Rothfels CJ, Johnson MG, Shaw AJ. 2016.** Analyses of transcriptome sequences reveal multiple ancient large-scale duplication events in the ancestor of Sphagnopsida (Bryophyta). *New Phytol* **211**(1): 300-318.

**[7] Bowman JL, Kohchi T, Yamato KT, Jenkins J, Shu S, Ishizaki K, Yamaoka S, Nishihama R, Nakamura Y, Berger F, et al. 2017.** Insights into Land Plant Evolution Garnered from the Marchantia polymorpha Genome. *Cell* **171**(2): 287-304 e215.

**[8] Banks JA, Nishiyama T, Hasebe M, Bowman JL, Gribskov M, dePamphilis C, Albert VA, Aono N, Aoyama T, Ambrose BA, et al. 2011.** The Selaginella Genome Identifies Genetic Changes Associated with the Evolution of Vascular Plants. *Science* **332**(6032): 960-963.

**Table S2 - Species contributions to single-copy orthologous groups**

| **Species** | **Single-copy Orthogroups** | **Percentage of all single-copy orthogroups** |
| --- | --- | --- |
| *Ceratodon purpureus* | 377 | 58.09 |
| *Syntrichia caninervis* | 499 | 76.89 |
| *Dicranum scoparium* | 521 | 80.28 |
| *Leucobryum glaucum* | 539 | 83.05 |
| *Scouleria aquatica* | 533 | 82.13 |
| *Thuidium delicatulum* | 600 | 92.45 |
| *Hypnum imponens* | 605 | 93.22 |
| *Anomodon rostratus* | 587 | 90.45 |
| *Climacium americanum* | 611 | 94.14 |
| *Plagiothecium laetum* | 581 | 89.52 |
| *Aulacomnium palustre* | 617 | 95.07 |
| *Rhodobryum ontariense* | 575 | 88.60 |
| *Bryum argenteum* | 346 | 53.31 |
| *Hedwigia ciliata* | 442 | 68.10 |
| *Timmia austriaca* | 422 | 65.02 |
| *Physcomitrella patens* | 649 | 100 |
| *Funaria Hygrometrica* | 597 | 91.99 |
| *Encalypta streptocarpa* | 373 | 57.47 |
| *Diphyscium foliosum* | 466 | 71.80 |
| *Buxbaumia aphylla* | 461 | 71.03 |
| *Atrichum angustatum* | 450 | 69.34 |
| *Polytrichum commune* | 421 | 64.87 |
| *Tetraphis pellucida* | 403 | 62.10 |
| *Andreaea rupestris* | 477 | 73.50 |
| *Eosphagnum inretortum* | 593 | 91.37 |
| *Sphagnum lescurii* | 409 | 63.02 |
| *Takakia lepidozioides* | 484 | 74.58 |
| *Marchantia polymorpha* | 649 | 100 |
| *Selaginella moellendorffii* | 635 | 97.84 |
